# Heme Impairs Alveolar Epithelial Sodium Channels Post Toxic Gas Inhalation

**DOI:** 10.1101/2020.01.22.909879

**Authors:** Saurabh Aggarwal, Ahmed Lazrak, Israr Ahmad, Zhihong Yu, Ayesha Bryant, James A. Mobley, David A. Ford, Sadis Matalon

## Abstract

We previously reported that cell-free heme (CFH) is increased in the plasma of patients with acute and chronic lung injury and causes pulmonary edema in animal model of acute respiratory distress syndrome (ARDS) post inhalation of halogen gas. However, the mechanisms by which CFH causes pulmonary edema are unclear. Herein we report for the first time the presence of CFH and chlorinated lipids (formed by the interaction of halogen gas, Cl_2_, with plasmalogens) in the plasma of patients and mice exposed to Cl_2_ gas. *Ex vivo* incubation of red blood cells (RBC) with halogenated lipids caused oxidative damage to RBC cytoskeletal protein spectrin, resulting in hemolysis and release of CFH. A single intramuscular injection of the heme-scavenging protein hemopexin (4 µg/kg body weight) in mice, one hour post halogen exposure, reversed RBC fragility and decreased CFH levels to those of air controls. Patch clamp and short circuit current measurements revealed that CFH inhibited the activity of amiloride-sensitive (ENaC) and cation sodium (Na^+^) channels in mouse alveolar cells and trans-epithelial Na^+^ transport across human airway cells with EC_50_ of 125 nM and 500 nM, respectively. Molecular modeling identified 22 putative heme-docking sites on ENaC (energy of binding range: 86-1563 kJ/mol) with at least 2 sites within its narrow transmembrane pore, potentially capable of blocking Na^+^ transport across the channel. In conclusion, results suggested that CFH mediated inhibition of ENaC activity may be responsible for pulmonary edema post inhalation injury.

## INTRODUCTION

Heme is an essential functional group of many proteins. However, non-encapsulated cell-free heme (CFH), a breakdown component of proteins such as hemoglobin, myoglobin, horseradish peroxidase, cytochrome *b*_5_, and cytochrome P450 can cause oxidative damage and impair cellular integrity (1) and is implicated in the pathogenesis of several disorders (2-10). CFH is an abundant source of redox-active iron capable of damaging lipid, protein, and DNA (11-13). In addition, CFH intercalates into cell membranes and disrupts plasma membrane integrity (14). Under normal conditions, circulating CFH is maintained at low levels (15) by serum albumin and hemopexin (16-19).

In a recent publication, Shaver et al found that cell-free hemoglobin levels in the air space correlated with alveolar-capillary barrier dysfunction in humans with acute respiratory distress syndrome (ARDS) (20). Further, they demonstrated that the intratracheal administration of cell-free hemoglobin to mice resulted in alveolar-capillary barrier disruption and acute lung injury (20). Interestingly, they also found that the effects of cell-free hemoglobin were mediated by the iron-containing heme moiety of cell-free hemoglobin, as intratracheal administration of free heme was sufficient to increase alveolar permeability in mice (20). In fact in our recent studies, we found that patients with chronic obstructive pulmonary disease (COPD) also had elevated levels of CFH (8) which correlated with the severity of the disease. In an animal model of ARDS, we found that scavenging CFH reduced pulmonary edema and improved lung function (9). However, the mechanisms of hemolysis and an increase in CFH in plasma in lung injury and ARDS are not known. Moreover, the mechanisms by which CFH impairs alveolar fluid clearance in ARDS is unclear.

The amiloride-sensitive epithelial Na^+^ channel (ENaC), an apical membrane protein complex, is the first and limiting step in Na^+^ transport across the alveolar space of humans and animals. This process is driven by the energy consuming Na^+^/K^+^-ATPase (sodium–potassium adenosine triphosphatase) located in the basolateral side of epithelial cells and plays a critical role in fluid reabsorption especially when the alveolar epithelium is damaged and permeability to plasma proteins is increased (21). Most important, this process of alveolar fluid clearance is impaired in ARDS (22). Ware et al. measured net alveolar fluid clearance in 79 patients with acute lung injury or ARDS and found that 56% had impaired alveolar fluid clearance, and 32% had submaximal clearance and only 13% had maximal clearance (22). They also found that patients with maximal alveolar fluid clearance had significantly lower mortality and a shorter duration of mechanical ventilation (22). A decline in Na^+^ and fluid clearance has also been reported in animal models of lung injury and ARDS. Vivona et al. demonstrated that hypoxia reduced Na^+^ conductance by attenuating ENaC activity (23). Polyubiquitination and decrease in cell surface stability of ENaC due to hypercapnia was reported by Gwozdzinska et al. as an important cause of decline in vectorial transport of Na^+^ across alveolar epithelium in an animal model of ARDS (24). In this manuscript, we explored whether CFH is responsible at least in part for ENaC impairment in ARDS.

We developed an animal model of ARDS by exposing C57BL/6 mice to halogen gases such as chlorine (Cl_2_) or bromine (Br_2_). We have previously shown that exposure to Br_2_ (600ppm, 30min) or Cl_2_ (400ppm, 30min) causes lung pathology similar to ARDS (9, 25-27) and that this injury is mediated at least in part by halogenated lipids, formed by the interaction of halogen gases with plasmalogens on the surface of lung epithelial cells, which are released into the alveolar spaces and plasma. Recent studies have shown that these halogenated lipids are elevated in patients who have ARDS (28) secondary to sepsis and also correlate with the disease severity (28).

In this study we measured for the first time the presence of chlorinated lipids and CFH in patients exposed to Cl_2_ gas during an industrial accident in 2019 at a water treatment facility in Birmingham, Alabama, US. We report that halogenated lipids increased oxidation of red blood cell (RBC) structural protein, spectrin, thereby increasing RBC fragility and rupture, resulting in the release of CFH in the plasma of mice exposed to halogen. To investigate the mechanisms responsible for the development of pulmonary edema, we evaluated whether CFH damages ENaC by patching lung epithelial cells and recording single channel activity in lung slices of mice exposed to halogens. To assess whether this injury was sufficient to decrease Na^+^ transport, we showed that CFH in concentrations found in alveolar space decreases short circuit current across confluent monolayers of human airway cells. Finally, our molecular modelling provides novel understandings by which heme inhibits ENaC. Therefore, we believe that is a first comprehensive study, which outlines the mechanisms of hemolysis and ENaC impairment in ARDS.

## RESULTS

### Cell-free heme (CFH) and chlorinated lipids are elevated in plasma of humans and animals exposed to Cl_2_ gas

Numerous studies including ours have previously shown that plasma levels of CFH are elevated in insults such as toxic halogen gas inhalation, sepsis, hyperoxia, and trauma, which ultimately leads to lung injury (2-10). However, the mechanisms of increased RBC fragility and plasma CFH are not known. We measured plasma CFH in 5 adult humans, which were admitted to the University of Alabama at Birmingham Emergency Department, post accidental exposure to Cl_2_ gas at the Birmingham water treatment plant. The average age of exposed humans was 48 years with 80% of them being males. Blood was also collected from corresponding age and sex matched non exposed humans. Plasma CFH levels were significantly increased in Cl_2_ gas exposed patients compared to their sex- and age- matched non-exposed individuals (Figure 1A). Person exposed to Cl_2_ gas also had elevated plasma levels of 16-carbon chlorinated fatty acid (ClFA) (Figure 1B) and 18-carbon ClFA (Figure 1C), as measured in plasma obtained from these individuals 3-6 h post exposure in the emergency room. These values were about 20 fold higher than those found in patients with sepsis (28). Similarly, exposure of adult C57BL/6 male mice to Cl_2_ gas (400ppm, 30 min) increased plasma levels of CFH levels (24 hours post exposure) (Figure 1D). In addition, we have previously shown that mice exposed to Cl_2_ gas also have elevated levels of chlorinated lipids (26).

**Figure 1.**
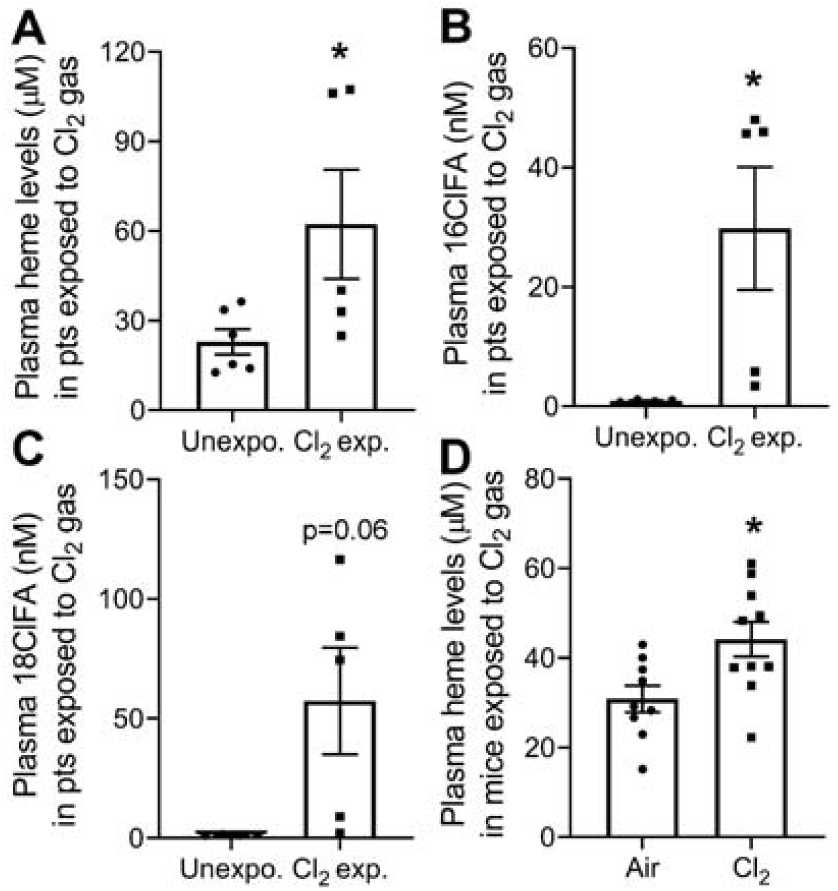
Plasma cell-free heme (CFH) and chlorinated lipids are elevated in humans and mice exposed to Cl_2_ gas. Blood was collected from patients exposed to Cl_2_ gas in the emergency room of the University of Alabama at Birmingham, 3-4 hours post exposure, stored for 72 hours at 4 °C at which time it was analyzed. Blood from human volunteers as controls was treated in the same fashion. Plasma CFH levels in persons exposed to Cl_2_ were higher than the age- and sex-matched human controls (n=5-6) (A). Cl_2_ exposed individuals also had elevated levels of 16ClFA (n=4-5) (B) and 18ClFA (n=4-5) (C). Similarly, adult male C57BL/6 mice exposed to Cl_2_ gas (400ppm, 30min) had increased levels of heme in plasma 24 hours post exposure (n=9-10) (D). Individual values and means ± SEM. \**P* < 0.05 vs. unexposed humans or air exposed mice; by unpaired t-test.

### Halogenated lipids increase RBC hemolysis and cell-free heme

To explore the role of halogenated lipids in increasing RBC fragility and plasma CFH levels, we first determined whether exposure to other halogen gases, such as bromine (Br_2_), also increases halogenated lipids. Male C57BL/6 mice were exposed to Br_2_ gas (600ppm, 30min) and then returned to room air. Using LC/MS quantitation, we found that the plasma levels of 16-carbon (Figure 2A-C) and 18-carbon (Figure 2D-F) free- and esterified-brominated fatty acids (BrFA) were elevated in the exposed animals. Even at 24 hours post exposure, the values of these variables were higher than those measured in patients with ARDS (28). We also identified elevated levels of brominated fatty aldehyde (BrFALD) in the broncholaveolar lavage fluid (BALF) (Supplementary Figure 1A, E) of exposed animals. Aldehydes can be oxidized to fatty acids which may exist either in the esterified (bound) or free form (26). The BALF levels of esterified- and free-BrFA were increased in mice exposed to Br_2_ (Supplementary Figure 1B-D, F-H). The aldehydes can also react with the antioxidant, glutathione, which exists in mM concentrations in the lung epithelial lining and plasma, to form glutathionylated fatty aldehyde (FALD-GSH) (26, 29, 30). We found high levels of 16- and 18- carbon FALD-GSH in the BALF (Supplementary Figure 2A-B), lung (Supplementary figure 2C-D), plasma (Supplementary Figure 3A-B), urine (Supplementary Figure 3C-D), and RBCs (Supplementary Figure 3E-F), of Br_2_ exposed mice. Because of their high reactivity, aldehydes could not be detected in the plasma in their natural configuration.

**Figure 2.**
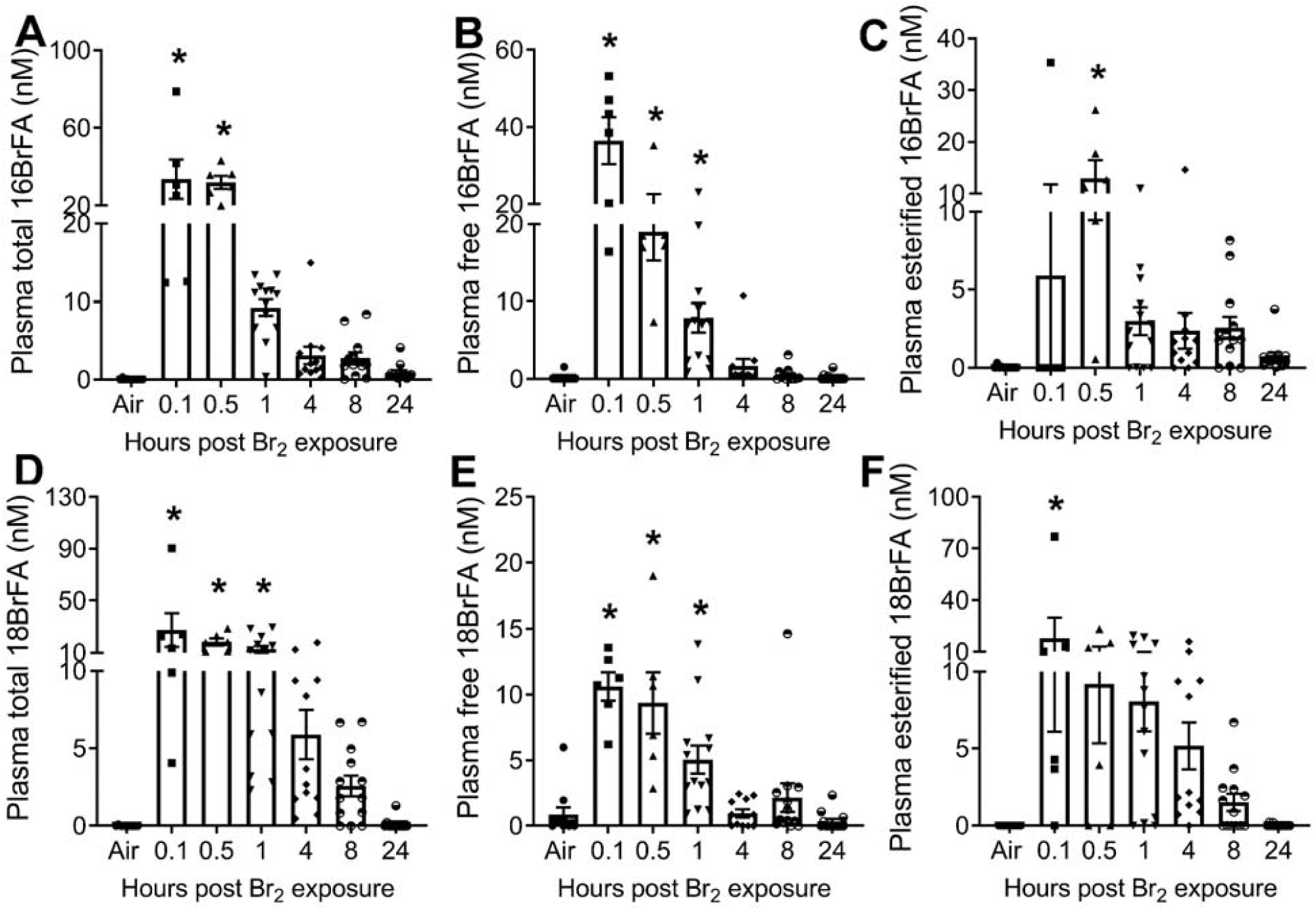
Brominated fatty acids (BrFA) are elevated in plasma of Br_2_ exposed mice. Adult male C57BL/6 mice were exposed to Br_2_ gas (600ppm, 30min) and returned to room air. Using ESI-LC/MS/MS quantitation, total, free, and esterified levels of 16- and 18- carbon BrFA were measured in plasma of mice at different intervals post exposure. The levels of total 16BrFA (n=6-13) (A), free 16BrFA (n=6-13) (B), and esterified 16BrFA (C) increased in plasma of Br_2_ exposed mice. Similarly, the levels of total 18BrFA (n=6-13) (D), free 18BrFA (n=6-13) (E), and esterified 18BrFA (F) increased in plasma of Br_2_ exposed mice. Individual values and means ± SEM. \**P* < 0.05 vs. air exposed mice, by one-way ANOVA followed by Tukey post hoc testing.

Next, to determine if halogenated lipids in the circulation or BALF are responsible for hemolysis and elevated CFH in plasma, RBCs were isolated from air exposed adult male C57BL/6 mice. The RBCs were then incubated *<u>ex vivo</u>* with the 16- and 18- carbon brominated or chlorinated lipids [fatty acid (FA) and fatty aldehyde (FALD), 1µM each], or their corresponding vehicle (FA and FALD) for 4 hours. The RBCs were then subjected to mechanical stress by mixing them with glass beads; the mixtures were shaken for 2 hours. Data showed that both the brominated and the chlorinated lipids increased the hemolysis of RBCs significantly (Figure 3A). In addition, the treatment of RBCs *<u>ex vivo</u>* with either brominated or chlorinated lipids resulted in increased oxidation of RBC membrane proteins, as indicated by elevated levels of carbonyl (aldehydes and ketones) adducts in protein side chains (Figure 3B-C).

**Figure 3.**
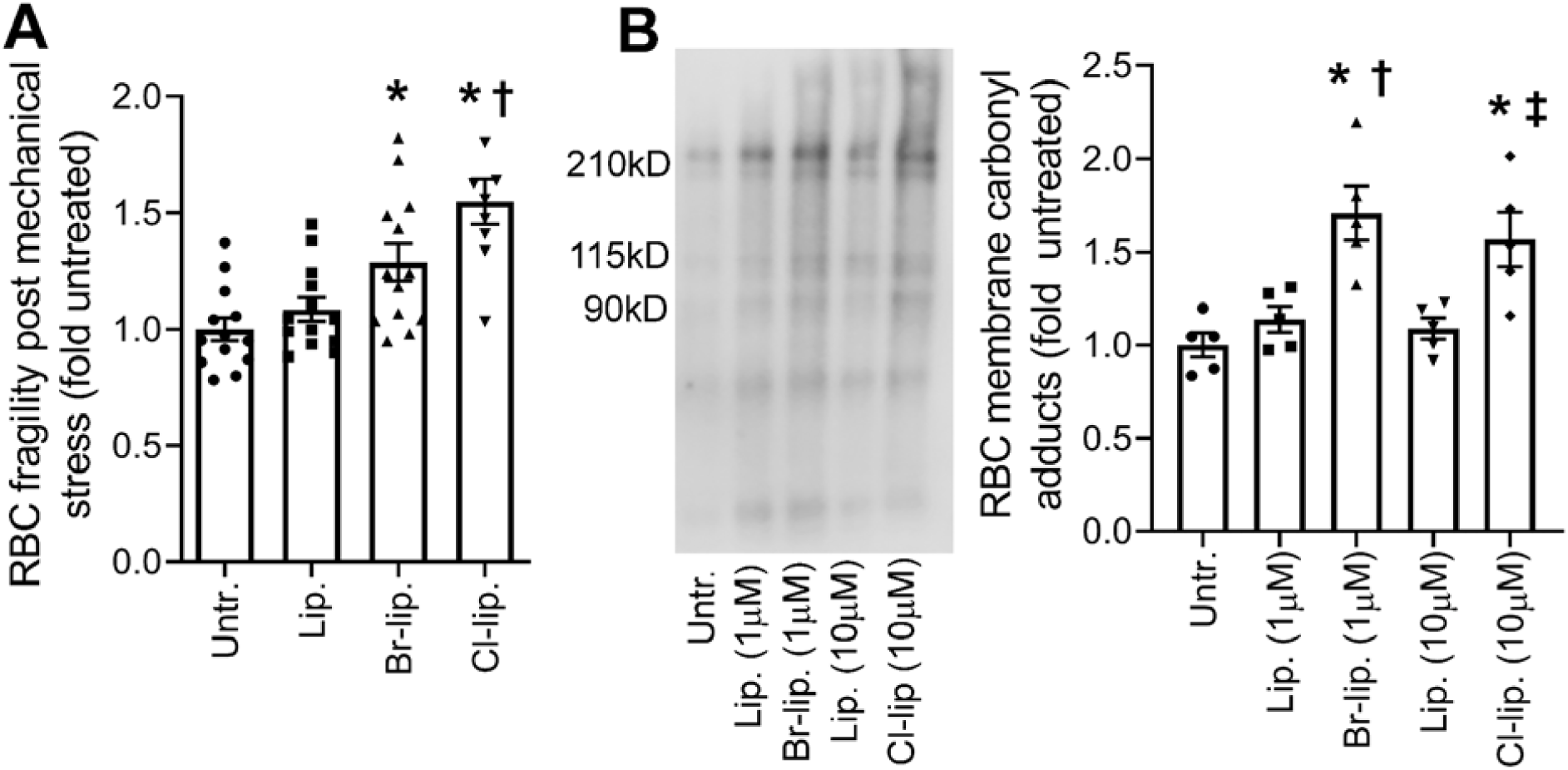
Halogenated lipids increase carbonylation and hemolysis of RBC. RBCs were isolated from adult male C57BL/6 mice and incubated *<u>ex vivo</u>* with Br-lip (16BrFA, 16BrFALD, 18BrFA, and 18BrFALD, 1µM each) (n=13), Cl-lip (16ClFA, 16ClFALD, 18ClFA, 18ClFALD, 1µM each) (n=9) in encapsulated liposomes, or equivalent amount of corresponding vehicle (16FA, 16FALD, 18FA, 18FALD, 1µM each) in encapsulated liposomes. After 4 hours incubation with these lipids, mechanical fragility of the cells was tested by mixing the RBCs with glass beads for 2 hours and then measuring the released hemoglobin as a result of hemolysis. Both brominated and chlorinated lipids increased RBC hemolysis (n=9) (A). In another set of experiments RBCs from adult male C57BL/6 mice were incubated *<u>ex vivo</u>* with the same composition of lipids with different concentrations and carbonyl adducts (aldehydes and ketones), a hallmark of the oxidation status of proteins, were measured by gel electrophoresis and western blotting using proteins obtained from the RBCs ghosts (n=5) (B). Both Br-lip and Cl-lip increased RBC carbonylation as indicated by increase in protein carbonyl adducts. Individual values and means ± SEM. \**P* < 0.05 vs. untreated RBCs, ^†^*P* < 0.05 versus non-halogenated lipids by one-way ANOVA followed by

To confirm the presence of carbonylation sites, as an indicator of oxidative stress and damage to RBC, we exposed adult male C57BL/6 mice to either Br_2_ (600ppm, 30min) or Cl_2_ (400ppm, 30min) gas. Twenty four hours post exposure, blood was drawn from these mice and RBC ghosts were isolated. High resolution LCMS2 identified carbonylation changes for 6 putative amino acid sites within the RBC structural protein, spectrin alpha chain, and one site within spectrin beta chain, 24 hours post-halogen exposure (Table 1). The sites listed were all confirmed using a number of high confidence filters that included A-score and localization probabilities as indicated in the table and in more detail within the methods section. While exposure to both Br_2_ and Cl_2_ induced carbonylation, Br_2_ appeared to have yielded a higher number of modifications indicating that it may be more damaging. Furthermore, each modification site appeared to be specific to the type of halogen gas, since no overlaps were found between the two exposures. The peptide marked with an asterisk in Table1 was found to be modified at K1988, and was chosen to be highlighted for the LCMS2 results (Figure 4), where an example of MS2 spectra are also illustrated to point out how these modifications can be putatively identified. Together, the results indicate that carbonylation of RBC structural proteins may induce hemolysis and increase CFH post exposure to toxic insult.

**Table 1.**
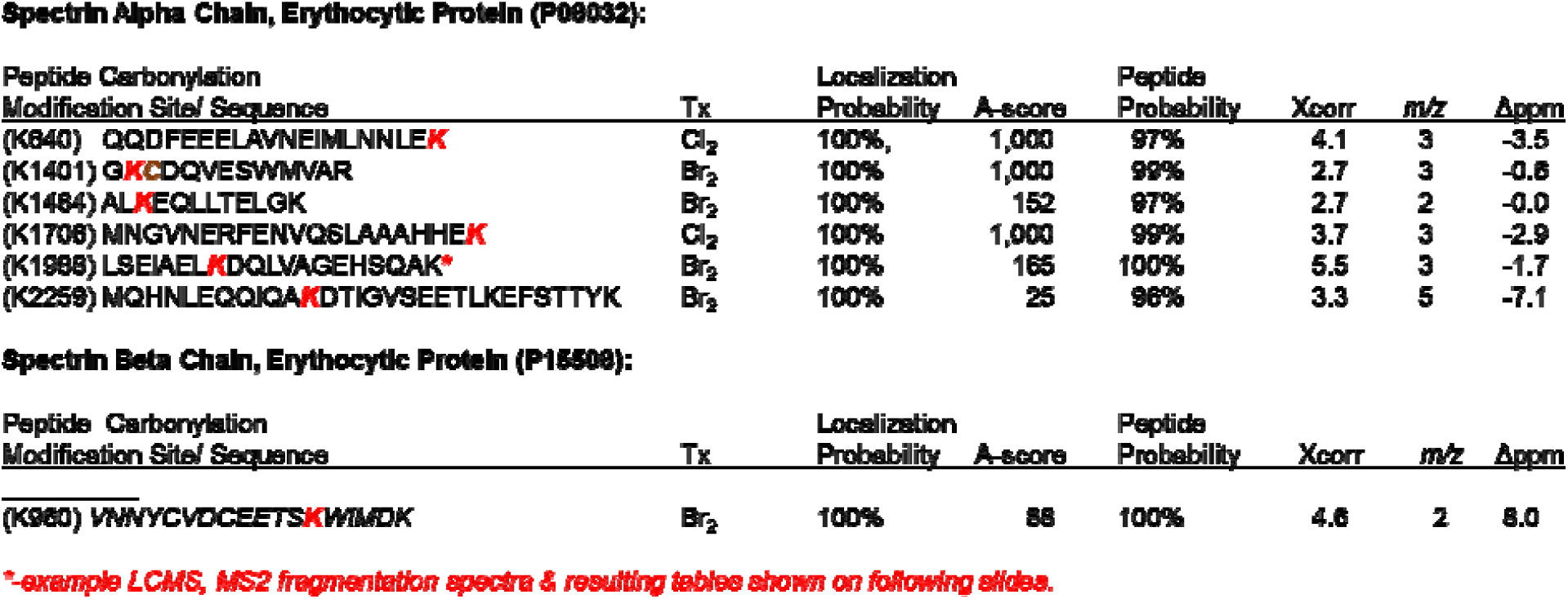
Carbonylation of spectrin alpha and beta chains from RBC plasma membranes isolated from mice 24 hours post-halogen exposure. Six putative carbonylation sites were confirmed by high resolution tandem mass spectrometry within spectrin alpha chain, and one site was identified within spectrin beta chain post-bromine exposure. These sites were all confirmed with a number of high confidence filters that included A-score and localization probabilities as indicated within the table, and further explained in detail within the methods section. The representative chromatography along with the MS1 & MS2 spectra are further highlighted in figure 4.

**Spectrin Alpha Chain, Erythrocytic Protein (P08032):**
| Peptide Carbonylation<br>Modification Site/ Sequence | Tx | Localization<br>Probability | A-score | Peptide<br>Probability | Xcorr | m/z | Appm |
| --- | --- | --- | --- | --- | --- | --- | --- |
| (K840) QQDFEELAVNEIMLNLEK | Cl<sub>2</sub> | 100% | 1,000 | 97% | 4.1 | 3 | -3.5 |
| (K1401) GKCDQVESWMMVAR | Br<sub>2</sub> | 100% | 1,000 | 99% | 2.7 | 3 | -0.8 |
| (K1484) ALKEQLLTELK | Br<sub>2</sub> | 100% | 152 | 97% | 2.7 | 2 | -0.0 |
| (K1708) MNGVNERFENVQSLAAHHEK | Cl<sub>2</sub> | 100% | 1,000 | 99% | 3.7 | 3 | -2.9 |
| (K1988) LSEIAELKDQLVAGEHSQAK* | Br<sub>2</sub> | 100% | 165 | 100% | 5.5 | 3 | -1.7 |
| (K2258) MQHNLEQQIQAKDTIGVSEETLKEFSTTYK | Br<sub>2</sub> | 100% | 25 | 96% | 3.3 | 5 | -7.1 |

**Table 1.** Carbonylation of spectrin alpha and beta chains from RBC plasma membranes isolated from mice 24 hours post-halogen exposure.
| Peptide Carbonylation<br>Modification Site/ Sequence | Tx | Localization<br>Probability | A-score | Peptide<br>Probability | Xcorr | m/z | Appm |
| --- | --- | --- | --- | --- | --- | --- | --- |
| (K980) VMNYCVDCEETSKWIMDK | Br<sub>2</sub> | 100% | 88 | 100% | 4.6 | 2 | 8.0 |

\*-example LCMS, MS2 fragmentation spectra & resulting tables shown on following slides.

**Figure 4.**
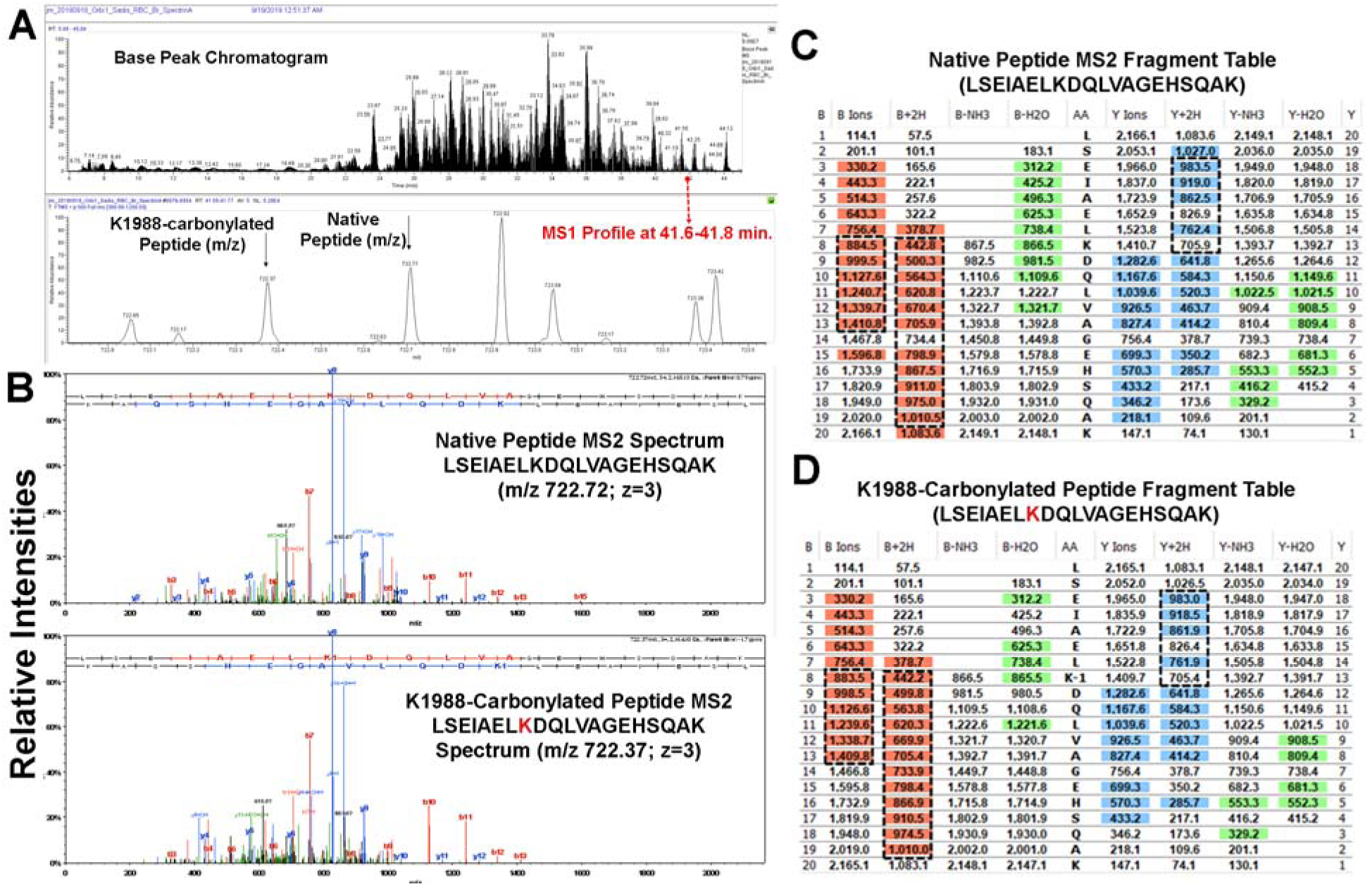
Halogen gas exposure increased carbonylation of RBC spectrin A and B chains in mice. A representative LCMS2 analysis is illustrated with a focus on the tryptic peptide (LSEIAELKDQLVAGEHSQAK) from spectrin alpha chain as highlighted in **Table 1**. (A) Illustration of the LCMS base-peak chromatogram, (B) parent-ion spectra from the peptide peaks of interest using data dependent analysis at 41.7 minutes along with the resulting MS2 spectra for those ions representing the native (top) vs. Ox-K1988 (bottom) peptide of interest. The tables, (C) native peptide, and (D) Ox-K1988 peptide represent the corresponding fragments for the peptides from the MS2 spectra. The highlighted sections (b-ions/ orange, y-ions/ blue) represent those fragments that were observed in the MS2 spectra. The MS2 spectra illustrate that the two peptides (native vs. modified) fragment are nearly identically at charge state 3, with only 1-dalton shifts that indicate K1988 is modified, which is made clear in the fragmentation tables. It’s noted that all the 1-dalton fragmentation shifts between the two tables are apparent below the b-8 ion and above the y-13 ion, both corresponding with AA#8 (K1988). MS was performed from RBC ghosts pooled from 4-5

### Heme scavenging attenuates RBC plasma membrane protein carbonylation and hemolysis

Next, we determined whether scavenging CFH would improve the integrity of RBC’s plasma membrane and prevent hemolysis. We exposed C57BL/6 mice to Br_2_ (600ppm) or Cl_2_ (400ppm) for 30 minutes and then treated the mice with an intramuscular injection of the heme scavenging protein, hemopexin, (4mg/kg BW) 1 hour later as published earlier (8, 9). RBCs were isolated from mice 1 day post exposure. Results demonstrated that hemopexin attenuated RBC plasma membrane protein carbonyl adducts post Br_2_ (Figure 5A) and Cl_2_ (Figure 5B) gas exposure. The exposure of isolated RBC to mechanical stress also showed that hemopexin reduced RBC hemolysis in the RBCs obtained from mice exposed to Br_2_ or Cl_2_ gas (Figure 5C). Together these results show that CFH released post exposure to halogen gas or halogenated lipids exaggerates RBC hemolysis and further contributes to RBC defect.

**Figure 5.**
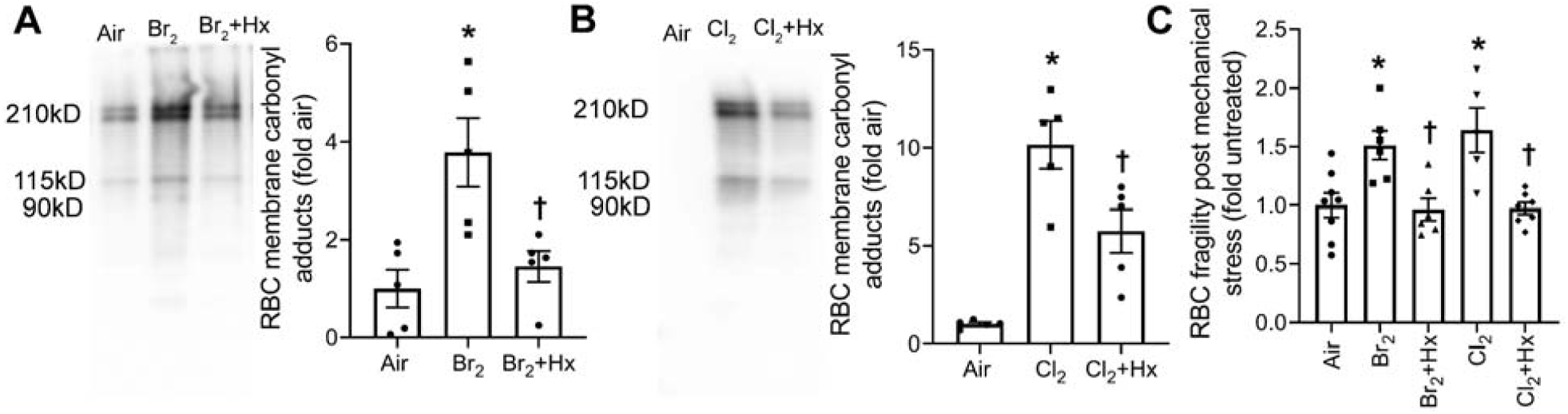
Heme scavenging attenuates RBC membrane protein oxidation and RBC fragility. Adult male C57BL/6 mice were exposed to Br_2_ (600ppm, 30min) or Cl_2_ (400ppm, 30min) gas and returned them to room air. One hour later, mice were given an intramuscular injection of either saline or hemopexin (Hx) (4mg/kg BW). Mice were sacrificed 24 hours post exposure and the RBCs were isolated. The carbonyl (aldehydes and ketones) adducts, a hallmark of the oxidation status of proteins, were measured by gel electrophoresis and western blotting using proteins obtained from the RBCs ghosts. The data demonstrated that Hx attenuated Br_2_ (n=5) (A) and Cl_2_ (n=5-6) (B) induced increase in RBC membrane protein oxidation. Further, subjecting the isolated RBCs to mechanical stress, showed that Hx treatment prevented the increase in RBC fragility and hemolysis induced by exposure of mice to the halogen gases (n=5-8) (C). Individual values and means ± SEM. \**P* < 0.05 vs. air exposed mice, ^†^*P* < 0.05 vs. mice exposed to Br_2_ for panel A, Cl_2_ for panel B, and their respective Br_2_ or Cl_2_ for panel C. The results were analyzed by one-way ANOVA followed by Tukey post hoc testing.

### Heme impairs ENaC activity and Na^+^ transport across lung epithelium

We have previously shown that CFH is responsible for lung edema and lung injury post toxic gas inhalation (8-10). Therefore, to determine the mechanism of CFH-induced lung edema, in the first series of experiments, we determined whether heme inhibits active Na^+^ transport across human bronchial epithelial cells (HBEC). Hemin (ferric chloride heme, mentioned just as heme throughout the text) was added in both the apical and basolateral compartments of Ussing chambers mounted with confluent monolayers as previously described (31, 32). Data demonstrated that heme inhibited amiloride-sensitive, ENaC, but not forskolin-stimulated, GlyH-101–inhibited, CFTR, currents within few seconds post exposure (Figure 6A-B) with an IC_50_ of about 500nM (Figure 6C). Transepithelial resistance did not decrease with increased heme concentrations but even at a heme concentration of 10 µM, remained above 1500 Ohms*cm^2^, indicating that the monolayers remained intact (data not shown).

**Figure 6.**
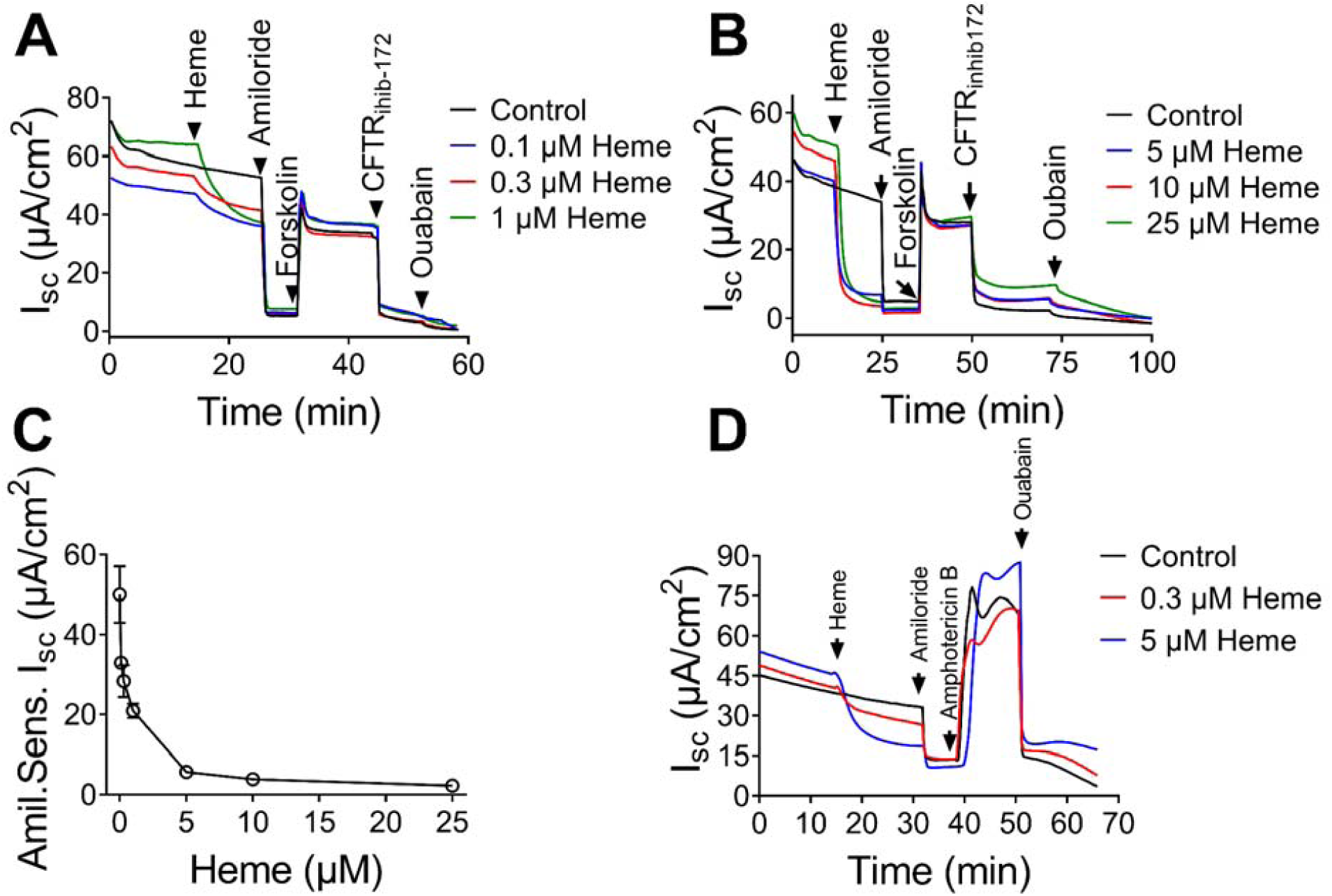
Heme impairs short circuit current (Isc) in human bronchiolar epithelial cells. Panels (A) and (B) illustrate the dose response of heme on short circuit current recorded from human bronchiolar epithelial cells monolayers mounted in Ussing chambers. Heme significantly reduced amiloride sensitive Na^+^ current in a dose-response fashion within seconds from its addition in the apical chamber but had no effect on the forskolin activated chloride current. The rate of Na^+^ current inhibition by heme is similar to Na^+^ current inhibition by amiloride (n=7). Panel (C) summarizes the dose response of Na^+^ current to increasing heme concentrations (n=7). Values are means ± 1 SEM. Further, cell monolayers were mounted on the Ussing system, Na^+^ current was inhibited by the addition of amiloride (10 µM) or heme (5 µM) in the apical compartment; then the apical membranes were permeabilized by the addition of amphotericin B in the apical compartment and after the current had stabilized, ouabain was added into the basolateral compartment. The difference in current prior to and following ouabain addition represents the Na^+^/K^+^ -ATPase (pump current). The data shows that heme does not inhibit the pump current (n=7). The graphs are representative of experiment which was repeated 7 times.

In the next series of experiments, we permeabilized the apical membranes of these monolayers and measured the ouabain-sensitive components of Na^+^/K^+^-ATPase activity. Data showed that heme did not impair (alter) Na^+^/K^+^-ATPase function even at concentrations that totally inhibited ENaC activity (5-25 µM) (Figure 6D). To further demonstrate that heme inhibited ENaC, we injected *Xenopus* oocytes with α-, β-, and γ-human ENaC cRNAs and measured current-voltage relationships 48 hours later. Oocytes injected with ENaC express significant amounts of Na^+^ currents, 90% of which are inhibited by 10μM amiloride (33). Treatment of oocytes with 1µM of heme immediately inhibited whole-cell Na^+^ current by 80% (Figure 7A-C). This decline in whole-cell Na^+^ current was similar to amiloride-mediated inhibition of Na^+^ current (Figure 7D-F).

**Figure 7.**
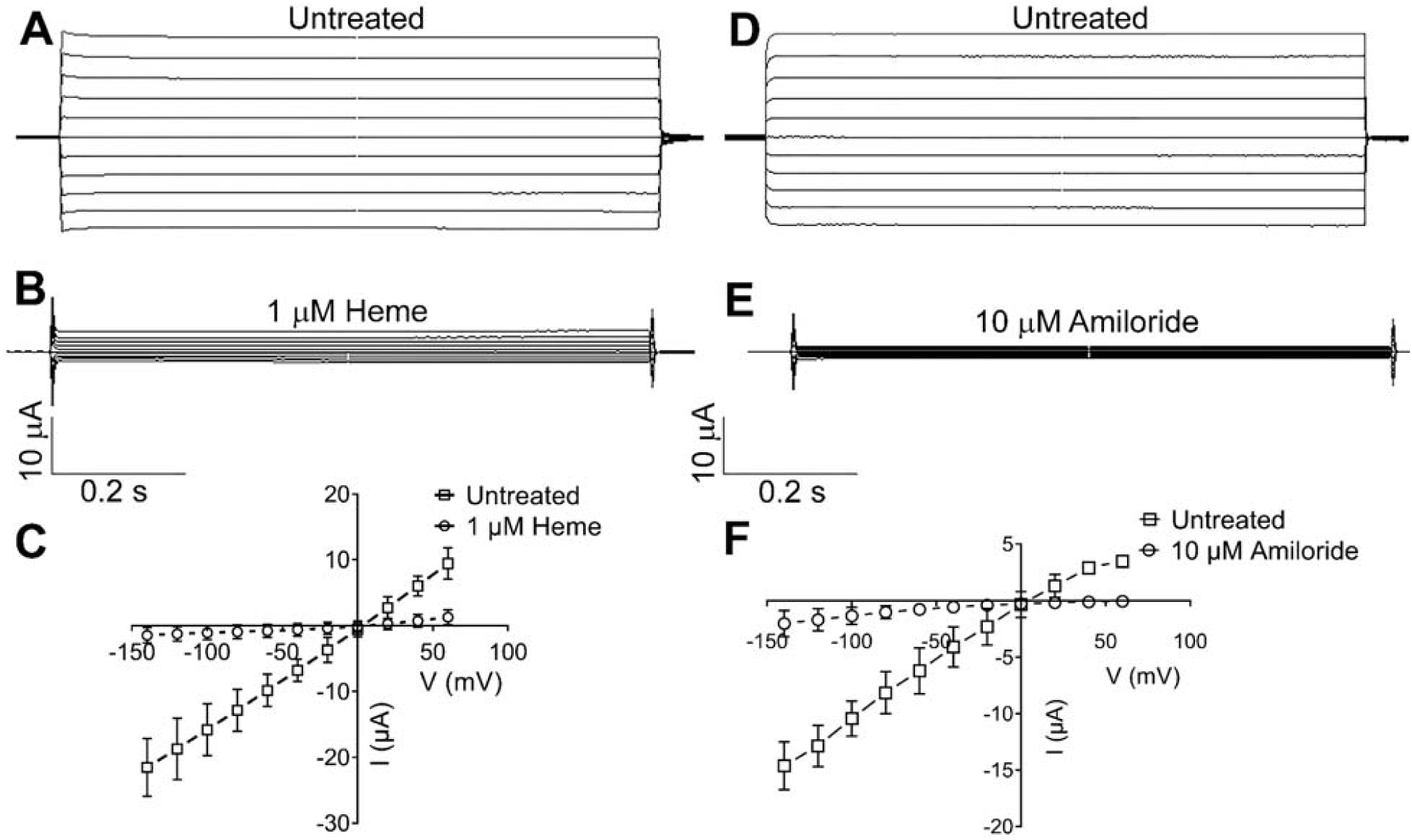
Heme inhibits ENaC heterologously expressed in oocytes. Panel (A) shows whole cell total current recording using the double voltage clamp technique from an oocyte 24 hours post injection of human –αβγ ENaC. Panel (B) shows that Na^+^ current is inhibited by 5µM heme in the bath. Panel (C) shows current-voltage relationships of total currents expressed in ENaC injected oocytes following perfusion with hemin or saline. (n = 11). Panels (D), (E), and (F) show a comparable inhibition of whole cell total current in oocytes by amiloride (n = 11). Values are means ± SEM.

Further, ENaC activity was measured in alveolar type II (AT2) cells in-situ (lung slices) using the cell-attached mode of patch-clamp technique as previously reported (33, 34). Heme was added in the upper portion of the pipette which allowed for the recording of baseline ENaC activity prior to heme reaching the membrane patch under the pipette. Recordings from AT2 cells show the activity of two characteristic conductances: a 4 pico siemens (pS) conductance of the highly Na^+^ selective channel (Figure 8A) and a 16 pS conductance of the non-selective cation channel (Figure 8E). However, once heme diffused through the pipette and reached the membrane patch under the pipette tip, it decreased the open probabilities of both the 4 pS (Figure 8B-C) and 16 pS channels (Figure 8F-G) with an IC_50_ of 125 nM for both 4 pS (Figure 8D) and 16 pS (Figure 8H).

**Figure 8.**
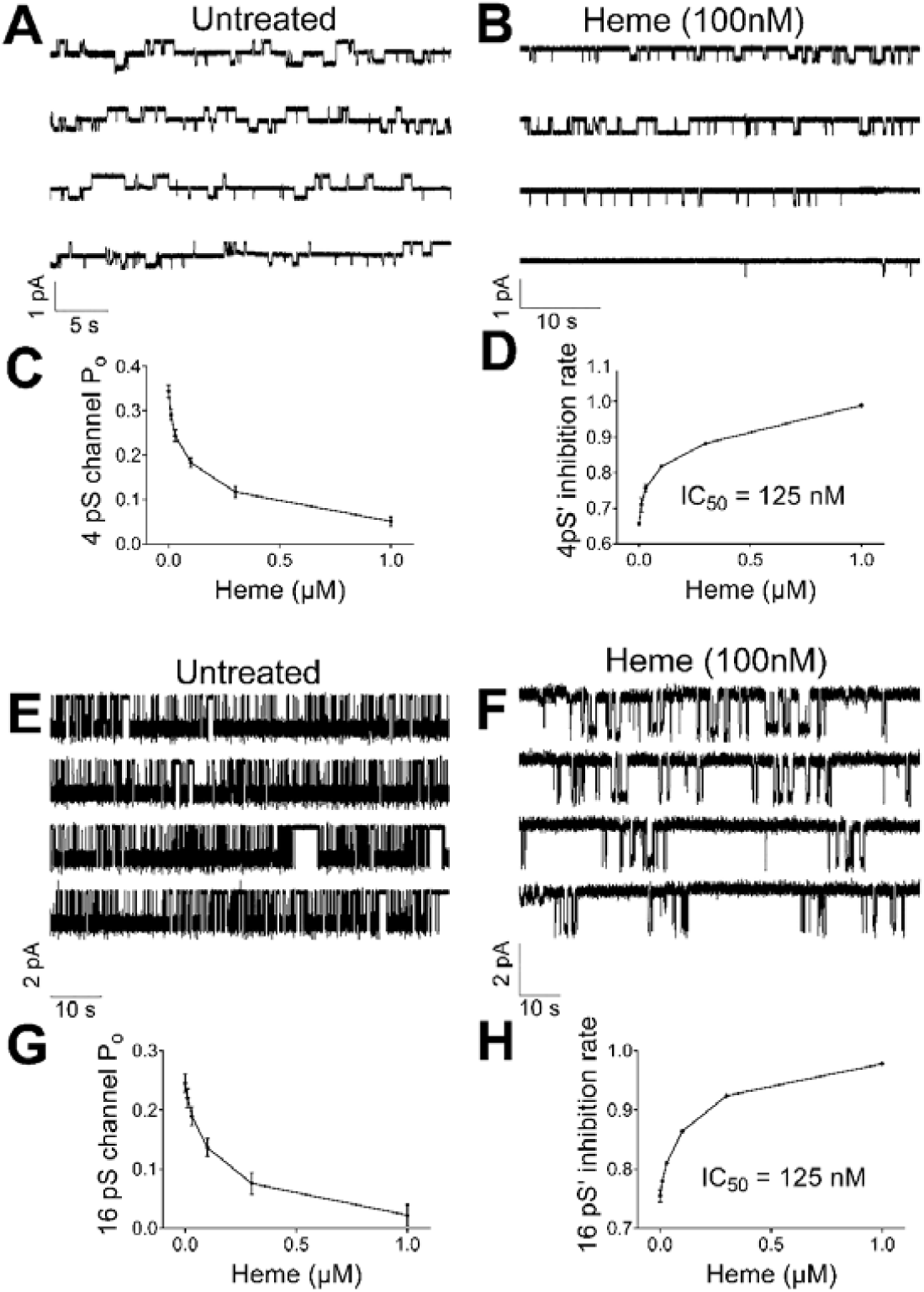
Heme inhibits ENaC activity in AT2 cells in situ. AT2 cells in lung slices were patched in the cell-attached mode. The top portion of the pipette was filled with heme so the overall concentration of heme in the pipette was 100 nM. Panel (A) represents a typical trace exhibiting mainly the 4 pS amiloride-sensitive current (ENaC). Notice the inhibition of ENaC activity as heme diffuses towards the patch under the pipette (B). Panel (C) shows the inhibition of open probability (P_0_) of 4 pS channel at increasing heme concentrations (0.01 to 1000 nM) in the pipette (n=9). Panel (D) illustrates the rate of ENaC inhibition by increasing heme concentrations: 10 nM, 30 nM, 100 nM, 300 nM and 1µM. The IC_50_ was 125 nM (n = 9). Panel (E) shows a record of AT2 channel activity in which the cation (16 pS) was very prominent. As in panel A, the top portion of the pipette was filled with heme so the overall concentration of heme in the pipette was 100 nM. Note the slow decrease of channel’s activity with time as heme reaches the membrane patch under the pipette (F). Panel (G) shows the open probability of 16 pS channel at increasing heme concentration in the pipette (same as above). Panel H shows that the IC_50_ was again 125 nM (n =10). Values

To further understand the mechanism by which CFH instantaneously inhibits EnaC activity and Na^+^ conductance across the lung epithelium, we performed computer modeling using YASARA software (35, 36) to identify potential heme binding sites and their potential ability to block Na^+^ conductance. For this purpose, we used a recently developed cryo-electron microscopy structure of ENaC (Protein Data Bank: 6BQN) (37). The ion channel has large extracellular domains and a narrow transmembrane pore and the α:β:γ subunits are arranged in a counter-clockwise manner in a 1:1:1 stoichiometry (37). The software predicted 22 potential docking sites of heme on ENaC with the energy of binding ranging from 86 to1563 kJ/mol. Close analysis of these docking sites showed that at least two heme-bonding sites are located within the EnaC transmembrane pore (energy of binding 390.5 and 313.2 kJ/mol) (Figure 9A-B), which can potentially block Na^+^ transport through the channel. Together, these results demonstrated that heme mediated decrease in ENaC activity may be responsible for lung edema during ARDS.

**Figure 9.**
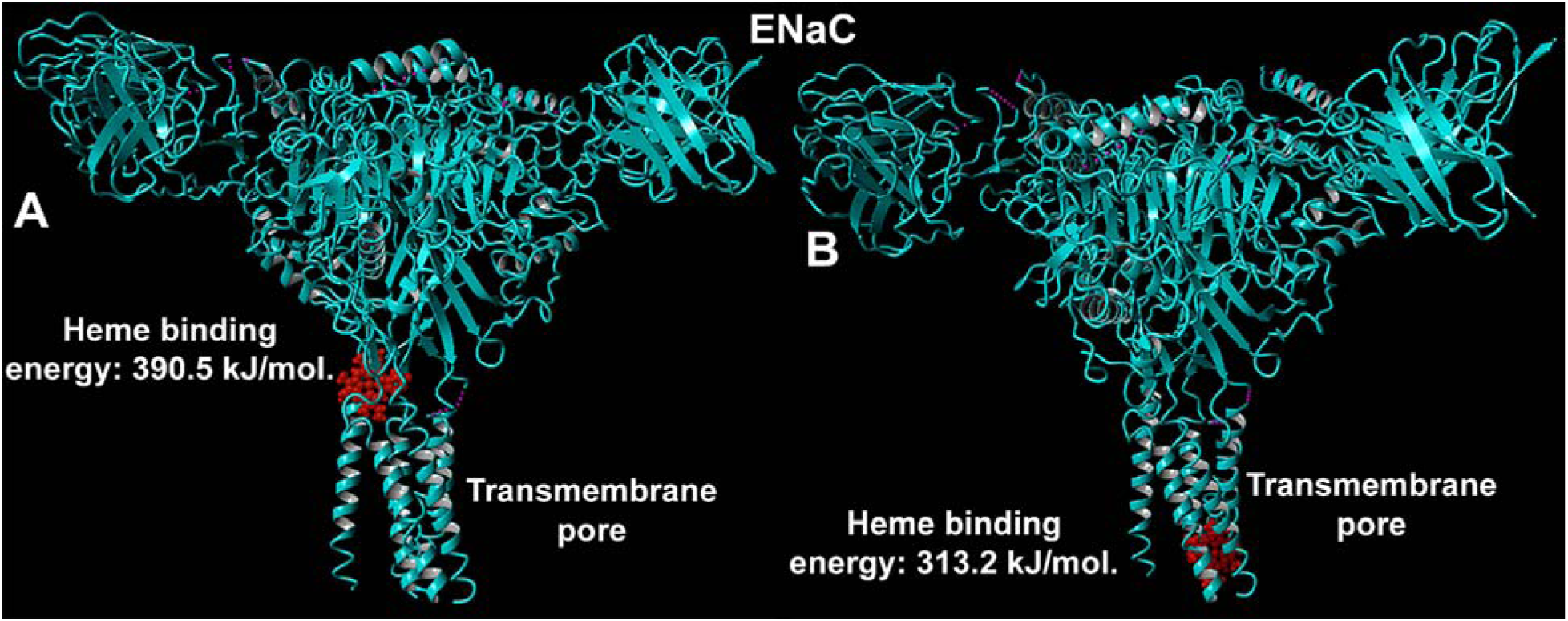
Computer modeling and heme docking on ENaC. The YASARA computer modeling software was used to identify heme binding sites on the known ENaC structure (Protein Data Bank: 6BQN). The AutoDock program was utilized to dock heme molecules on ENaC. The software identified 22 heme docking sites on ENaC. Two prominent heme docking sites within the transmembrane domain are highlighted in the figure. The energy of heme binding to ENaC at these two sites was determined by the software as 390.5kJ/mol (A) and 313.2 kJ/mol.

## DISCUSSION

ARDS is a common and severely morbid syndrome with high mortality despite years of advances in the understanding and management of this complex illness. Although several scientific theories on the pathogenesis of ARDS have developed in animals, they are often difficult to test in humans due to many clinical variables. We have previously established an animal model of inhalation lung injury and ARDS, where C57BL/6 mice were exposed to halogen gases such as Cl_2_, Br_2_, or phosgene (COCl_2_) (8-10, 38, 39). One day post exposure, rodents exposed to these agents developed lung pathology similar to ARDS patients such as accumulation of protein rich fluid in alveoli, migration of peripheral immune cells in the lung, and release of cytokines and chemokines in lung and plasma; a significant portion died within the first 48 hours post exposure from respiratory failure (9, 10). Those which survived developed peribronchial fibrosis and pulmonary emphysema (8). Interestingly, we also found that the rodents exposed to toxic gases such as Br_2_ or COCl_2_ had elevated plasma and BALF levels of CFH (8-10). The administration of the heme scavenging protein, hemopexin, or the overexpression of the heme degrading enzyme, heme-oxygenase 1 (HO-1), in Br_2_ exposed animals resulted in significant reduction of lung edema and inflammation (8, 9).

Hemolysis has been reported in several clinical conditions (2-10). Under hemolytic states, the physiological mechanisms for removing CFH from the circulation or heme detoxification systems get overwhelmed, which allows for nonspecific heme uptake and heme catalyzed oxidation reactions (40-42). In this manuscript we report for the first time, an increase in the plasma levels of CFH in humans who were exposed to Cl_2_ gas post accidental gas leakage at the Birmingham Water Works Plant on February 27^th^, 2019. These findings correlated with our rodent experiments, where we found that C57BL/6 mice exposed to Cl_2_ gas also had elevated plasma levels of CFH. Since halogen gases such as Cl_2_ and Br_2_ do not cross alveolar epithelium post inhalation, to reach blood stream, they are unlikely to cause hemolysis. However, we had earlier reported that rodents exposed to Cl_2_ gas had elevated plasma levels of chlorinated fatty acids (ClFA), which are formed by the interaction of Cl_2_ with plasmalogens on the alveolar epithelium (26). Therefore, we sought to determine whether these longer lasting, modified lipids, are the catalyst for hemolysis and elevated CFH in animals and humans. *Ex vivo* exposure of RBCs to chlorinated or brominated lipids in concentrations similar to seen in animals post halogen gas exposure resulted in hemolysis and release of CFH. Interestingly, we found that ClFA were also elevated in humans that were accidently exposed to Cl_2_ gas suggesting that halogenated lipids may be responsible for hemolysis post inhalation of toxic gases. Moreover, recent studies have shown that these halogenated lipids are elevated in patients who have ARDS secondary to sepsis and correlate with the disease severity (28). In these patients, chlorinated and brominated lipids are increased as an end product of reactions by myeloperoxidase (MPO) and eosinophil peroxidase (EPO) respectively (43). Thus, our data establishes that halogenated lipids are both biomarkers and potential mediators of hemolysis and acute lung injury.

Additionally, an increase in plasma CFH can itself cause substantial hemolysis by stimulating potassium loss and impairing the ability of RBCs to maintain cation gradient (44). CFH also disrupt lipid–lipid, lipid–protein, and protein–protein associations in RBC ghosts (45). Earlier studies have shown CFH can destabilize RBC membrane by altering the conformation of cytoskeletal proteins, such as spectrin and protein 4.1 (46). The hydrophobic molecule, CFH, enters RBC membrane and interacts with hydrogen peroxide to cause oxidative modification of cytoskeletal proteins (47). Oxidation of amino acid residues in spectrin can result in the aggregation of the protein, which may cause a decrease in spectrin content on the RBC membrane (48). In the present study, we found that the exposure to Cl_2_ or Br_2_ gas resulted in the post translational modification (carbonylation, a read out of oxidative stress) of the lysine residues in spectrin alpha and beta chains. Interestingly, we report for the first time that Cl_2_ and Br_2_ gases caused carbonylation of distinct lysine residues in spectrin. Specifically, Br_2_ was able to cause carbonylation of both alpha and beta chains of spectrin, while Cl_2_ exposure caused the carbonylation in spectrin alpha chain only. These results suggest that the initial hemolysis and release of CFH by halogenated lipids can potentially be self-propagated by CFH in ARDS resulting in continued hemolysis even when the initial insult ceases to exist.

Recent studies have shown that CFH disrupts alveolar-capillary barrier (20). A functional alveolar epithelial barrier is vital for optimal fluid balance and gas exchange in the lung (49, 50). Excess alveolar fluid from the air space is reabsorbed into the interstitium by active Na^+^ transport process in which Na^+^ enters the alveolar epithelial cells through the apically located ENaC and cation channels and is pumped out subsequently by the basolaterally located Na^+^/K^+^-ATPase (49, 51). Mice lacking ENaC are unable to clear lung fluid from the alveoli and die immediately after birth (52). Importantly, in most ARDS patients, alveolar fluid clearance (AFC) is impaired (22). Moreover, the mortality is significant higher in ARDS patients with impaired AFC than in ARDS patients with normal AFC (22) suggesting that ENaC dysfunction is critical in ARDS pathology. In addition, increased concentration of reactive intermediates following various forms of lung injury have been shown to also inhibit the Na^+^/K^+^-ATPase which abolishes active Na^+^ transport leading to alveolar edema (53-57). However, data presented herein clearly establish that in our models of lung injury, Na^+^/K^+^-ATPase is not inhibited and the decrease in short circuit current across cultured human airway cells post heme exposure is due to damage to the apically located Na^+^ conducting pathways, including ENaC.

ENaC is composed of 3 homologous subunits α, β and γ (58, 59). At the single channel level, ENaC is highly selective for Na^+^ over K^+^ with low conductance (∼4pS) with prolonged channel opening. ENaC activity is blocked by the potassium-sparing diuretic, amiloride, with a K_i_ of around 100nM (60). ENaC activity can be altered significantly in two ways. Firstly, a rapid alteration in Na^+^ conduction through ENaC can be achieved by changing the open probability (P_o_) or open state of the channel (61). The decline in open probability results in decreased Na^+^ transport without a change in the surface expression of the channel. Secondly, Na^+^ reabsorption through ENaC can be changed by altering the number of channels located in the apical membrane (62-64). In the present study, we found that heme rapidly (within few seconds) reduced the open probability of the 4 and 16 pS channels with an IC_50_ of 125 nM in mouse lung slices. However, the activity of the Na^+^/K^+^-ATPase or the cyclic AMP (cAMP)-activated chloride (Cl^-^) channel, CFTR, was not altered by heme exposure. In a different study, heme has been shown to reduce the open probability of ENaC in mouse kidney cortical-collecting duct cells with an IC_50_ of 23.3nM (65). The difference in the IC_50_ seen in our study could be explained by the different techniques used in each study. We applied heme to the outer side of the membrane, cell attached mode, while Wang et al. (65) applied heme to the inner side of the membrane using the inside-out configuration of the patch clamp. It is likely that the site to which heme binds ENaC and blocks the channel is closer to the inner narrower side of the transmembrane pore as shown in our model of putative heme binding sites on ENaC. In addition, the response of heme to particular cell and tissue type may be different since we used lung tissues and the other study used cultured kidney cells (65).

There are no known heme binding sites on ENaC. Therefore, to identify these potential sites of ENaC, we performed computer modeling using a known cryo-electron microscopy structure of ENaC (37). The software predicted 22 potential docking sites of heme molecule on ENaC with the energy of binding ranging from 86 to1563 kJ/mol. In particular, we found that at least two these heme-bonding sites were located near both ends of the narrow transmembrane pore of ENaC (energy of binding 390.5 and 313.2 kJ/mol). Therefore, it is entirely possible that the relative large molecular mass of the heme molecule (616.5 Da) blocks the pore of the channel and prevents the transportation of a smaller ion such as Na^+^ (22.9 Da). This would explain the immediate response of heme to Na^+^ transport. Previous studies on BK (calcium-activated potassium) channels have demonstrated that the CXXCH amino acid sequence serves as a conserved heme-binding motif (66). However, no individual ENaC subunit contains this motif and therefore it is possible that the combined allosteric structure of the 3 ENaC subunits provide basis for heme interaction. Further investigation involving truncation, replacement, or mutation of regions within the subunits will be required to fully understand this phenomenon. Since in our study, heme did not alter the activity of the Na^+^/K^+^-ATPase or the cyclic AMP (cAMP)-activated chloride (Cl^-^) channel, it also signifies that the inhibition by heme is channel specific. In conclusion, our study has provided a potential mechanism by which heme impairs fluid clearance and contributes to lung edema in ARDS.

## METHODOLOGY

### Human

The study was approved by the University of Alabama at Birmingham Institutional Review Board (IRB Protocol 300002065 and 300000860) and the Saint Louis University Institutional Review Board (IRB 9952). Demographic information were recorded on all volunteers and blood was drawn from peripheral vein. Plasma was isolated from the blood, aliquoted, and stored at −80°C using Freezerworks Sample Inventory Management software (Dataworks Development, Inc, Mountlake Terrace, WA, USA). No samples had undergone freeze-thaw cycles prior to use in the study.

### Animals

Adult male C57BL/6 mice (20-25g) were bought from Charles River, non-Frederick/NCI. All mice used in the study were males. All mice were raised under a 12-hour dim light/12-hour dark cycle with access to a standard diet and tap water ad libitum. Euthanasia protocol based on intraperitoneal injections of ketamine and xylazine was used in the study for mice to minimize pain and distress. All animal care and experimental procedures were approved by the Institutional Animal Care and Use Committee at the University of Alabama in Birmingham.

### Exposure to halogen gas

Mice were exposed to Br_2_ gas (600 ppm) or Cl_2_ gas (400ppm) in a cylindrical glass chamber for 30 minutes, as previously described (9, 26). Control mice were exposed to room air in the same experimental conditions as Br_2_ or Cl_2_ exposed mice. Exposures were performed with two mice in the same chamber at any one time, and all exposures were performed between 6:00 AM and 12:00 PM. Tanks were replaced when the pressure in the tanks reached 500 psi. In each case, immediately following exposure, mice were returned to room air. All experiments involving animals were conducted according to protocols approved by the UAB IACUC.

### Treatment of animals with hemopexin

Adult male C57BL/6 mice were exposed to Br_2_ gas (600 ppm), Cl_2_ gas (400ppm), or air in a cylindrical glass chamber for 30 minutes, as described above. Following exposure, mice were returned to room air and then 1 hour later, mice were treated with an intramuscular injection of either saline or purified human hemopexin (4mg/kg body weight, dissolved in saline) (Product No. 16-16-080513-LEL; Athens Research and Technology, Athens, GA). All experiments involving animals were conducted according to protocols approved by the UAB IACUC.

### Measurement of Br-lip

Mice were euthanized at various times post-Br_2_ gas exposure using a mixture of ketamine/xylazine (200/10 mg/kg) administered by intraperitoneal injection. Blood was collected via cardiac puncture, lungs were excised, and urine was collected from the bladder. Blood was centrifuged at 6,000 rpm for 5 min to obtain the plasma fraction. Lungs, urine, and plasma samples were flash-frozen in liquid nitrogen and stored at −80°C. In some mice, the lungs were lavaged, and the recovered bronchoalveolar lavage fluid (BALF) was centrifuged immediately at 3,000 *g* for 10 minutes to pellet the cells. Supernatants were flash-frozen. All samples were shipped overnight on dry ice to Dr. Ford at St. Louis University. Br-FALD was measured following conversion to its pentafluorobenzyl oxime using negative ion-chemical ionization GC/MS as previously described (67). Free, esterified, and total (free + esterified) Br-FA were measured as previously described for chlorine by LC/MS following Dole extraction (26, 68). Total lipids were measured by LC/MS after base hydrolysis and esterified Br-FA calculated by subtracting free lipids from total lipids. Extractions were performed using 25 µl of plasma spiked with 517 fmol of 2-chloro-[*d*_*4*_-7,7,8,8] palmitic acid (2-[*d*_*4*_]ClPA) as the internal standard, and for lungs, 40–50 mg of tissue was used, spiked with 20 pmol of 2-[*d*_*4*_]ClPA internal standard as mentioned earlier for Cl-FAs (26).

### Measurement of glutathione adducts of 2-Br-PALD

Plasma, lung, BALF, and urine samples were analyzed as previously described (26). Briefly, 25 µl of plasma, RBCs (diluted with 75 µl of water), BALF or urine were spiked with 90 fmol of [*d*_*4*_]HDAGSH and 10 mg of pulverized lung tissue was spiked with 900 fmol [*d*_*4*_]HDAGSH. Plasma, lung, RBCs, BALF, and urine were then extracted according to a similar Bligh and Dyer method as described for the Cl-lipids (68); however, the aqueous layer was saved as the GSH adducts partition to the aqueous layer. The organic layer was subsequently washed with 1 volume of methanol:water (1:1 v:v) and combined with the previous aqueous layer. The combined aqueous layers were diluted with 1/3 vol of water and extracted on a Strata-X followed by ESI-LC/MS/MS quantitation, as previously described (29).

### *Ex Vivo* RBC mechanical fragility

Blood was obtained from adult C57BL/6 mice in the presence of an anticoagulant and incubated with 1µM each of Br-lip (16BrFA, 16BrFALD, 18BrFA, 18BrFALD), Cl-lip (16ClFA, 16ClFALD, 18ClFA, or 18ClFALD) or the corresponding non-halogenated lipids as vehicle (16 and 18 carbon palmitic acid or palmitaldehyde) for 4 hours with rotations. In a separate set of experiments, blood was obtained from mice exposed to Br_2_, Cl_2_, or air in the presence or absence of treatment with hemopexin as mentioned above. Plasma was separated and the RBCs were washed with isotonic solution 3 times to remove traces of plasma. RBCs were then re-suspended in normal saline. The RBC suspensions along with 4×4mm glass beads (Pyrex) in DPBS were then rotated 360° for 2 hours at 24 rpm at 37°C. The RBC suspension was then centrifuged at 13,400g for 4 min to separate the intact or damaged cells from the supernatant containing heme/hemoglobin from the lysed cells during this mechanical stress. Free heme/hemoglobin was transferred into a new tube and the absorbance of the supernatant recorded at 540nm as described earlier (69). Subsequently, one hundred percent hemolysis of RBCs was achieved by treating them with 1% Triton x-100 solution. The fractional hemolysis of the sample was then obtained by dividing the optical density of the sample by the optical density of the 100% hemolyzed sample.

### Measurement of protein carbonyl adducts in RBC ghosts

RBCs were separated from the plasma and hemolyzed with 20 mM hypotonic Hepes Buffer. The mixture was centrifuged at 14000 g for 20 min and RBC pellet was dissolved in RIPA buffer (Thermo Fisher Scientific, MA). The protein was quantified by the BCA method and equal amounts of proteins (10µg) were loaded into a 4-20% gradient gel and proteins were separated and stained with Amido Black (Sigma-Aldrich, St Louis, MS). The presence of protein carbonyl adducts in RBC ghosts were assessed using the Oxyblot protein oxidation detection kit (Product number: S7150, EMD Millipore, Billerica, MA), according to the manufacturer’s protocol. Briefly, the carbonyl groups in the protein side chains were derivatized to 2,4-dinitrophenylhydrazone by reacting with 2,4-dinitrophenylhydrazine. Precisely, 10 µg of protein was used for each sample, and the 2,4-dinitrophenol-derivatized protein samples were separated by polyacrylamide gel electrophoresis, as described previously (9). Polyvinylidene fluoride membranes were incubated for 1 hour in the stock primary antibody (1:150 in 1% PBS/TBST buffer), and after washing, for 1 hour in the stock secondary antibody (1:300 in % PBS/TBST buffer). Membranes were washed 3× in TBST and visualized. The abundance of protein carbonylation was assessed by densitometry of each lane and normalization for each lane protein loading was done by SDS PAGE gel quantification.

### Lung slices preparation

Eight-week-old C57BL/6 male mice (∼20–25 g body weight) were purchased from The Jackson Laboratory (Bar Harbor, ME). Lung slices were prepared as previously described (70). The right lower lobes were dissected, attached to tissue holder using cyanoacrylate adhesive gel, and sectioned into slices of 200 μm thick. The slices were transferred to a six-well plate containing Dulbecco’s Modified Eagle’s Medium without serum, supplemented with penicillin–streptomycin, and allowed to recover at 37°C in a humidified environment of 95% air/5% CO_2_ for 2 to 3 hours.

### ENaC single channel activity in AT2 cells *in situ*

A lung slice was transferred to the recording chamber on the stage of an upright Olympus microscope EX51WI (Olympus, Pittsburgh, PA). Single-channel activity in AT2 cells was recorded using the cell-attached mode of the patch clamp technique (33, 71). AT2 cells were identified by the presence of scattered green fluorescence after incubation with Lysotracker Green (catalogue number DND-26; Invitrogen, Eugene, OR).

### Human bronchial epithelial cells isolation, culture, and short circuit currents recording

Human bronchiolar epithelial cells (HBECs) were provided by UAB CF Center upon request. Cells were isolated from human lungs not used for transplantation, HBECs were seeded onto permeable support and allowed to form confluent monolayers (3 to 4 weeks in culture). Tight monolayers were mounted in an Ussing chambers system and short circuit currents were monitored and recorded using an amplifier (physiologic instruments, San Diego, CA). Monolayers were bathed in Ringer solution bubbled with 95% air 5% Co_2._ Hemin (ferric chloride heme, mentioned as heme throughout the text), dissolved in DMSO, was added to both sides of monolayers.

### ENaC expression in Xenopus oocytes

Detailed description of these techniques has been previously reported (71). In brief, oocytes isolated from *Xenopus laevis* frogs, were injected with cRNAs encoding for wild-type α, β, γ-hENaC (8.4 ng each), dissolved in 50 nl of RNase-free water per oocyte, and incubated in half-strength L-15 medium for 24 to 48 hours. Whole-cell cation currents were measured by the two-electrode voltage clamp. A TEV 200 voltage clamp amplifier (Dagan Corp., Minneapolis, MN) was used hold oocytes membrane potential at −40 mV. Current–voltage (I–V) relationships were obtained by stepping the Vm from −140 mV to +60 mV in 20 mV increments. Sampling protocols were generated by pCLAMP 9.0 (Molecular Devices, Union City, CA). Currents were sampled at the rate of 1 kHz, filtered at 500 Hz, and simultaneously stored electronically and displayed in real time. Hemin was diluted to the desired final concentration in ND96 and applied to the oocytes through the perfusion system at a rate of 1 ml/ min.

### Mass spectrometry

#### Sample Preparation

*(1D separations was carried out by SM group, may need to be modified)* Samples were denatured in 1xfinal NuPAGE™ LDS Sample Buffer (Cat.#NP0007, Invitrogen), and the resultant enriched proteins were separated onto a NuPAGE™ 10% Bis-Tris Protein gel (Cat.# NP0315BOX, Invitrogen) at 200V constant for 25min. The gel was stained using a Colloidal Blue Staining Kit (Cat.#LC6025, Invitrogen) following manufacturer’s instruction. Each gel lane was excised (two equal-sized fractions pertaining to spectrin alpha and beta molecular weights) and digested overnight at 37°C with Pierce™ Trypsin Protease, MS Grade (Cat.#90058, Thermo Scientific) as per manufacturer’s instruction. Digests were reconstituted in 0.1%FA in 5:95 ACN:ddH2O at ∼0.1ug/uL.

#### nLC-ESI-MS2 Analysis & Database Searches

Peptide digests (8µL each) were injected onto a 1260 Infinity nHPLC stack (Agilent Technologies), and separated using a 100 micron I.D. x 13.5 cm pulled tip C-18 column (Jupiter C-18 300 Å, 5 micron, Phenomenex). This system runs in-line with a Thermo Orbitrap Velos Pro hybrid mass spectrometer, equipped with a nano-electrospray source (Thermo Fisher Scientific), and all data were collected in CID mode. The nHPLC was configured with binary mobile phases that included solvent A (0.1%FA in ddH2O), and solvent B (0.1%FA in 15% ddH2O / 85% ACN), programmed as follows; 10min @ 5%B (2µL/ min, load), 60min @ 5%-40%B (linear: 0.5nL/ min, analyze), 5min @ 70%B (2µL/ min, wash), 10min @ 0%B (2µL/ min, equilibrate). Following each parent ion scan (300-1200m/z @ 60k resolution), fragmentation data (MS2) was collected on the top most intense 15 ions. For data dependent scans, charge state screening and dynamic exclusion were enabled with a repeat count of 2, repeat duration of 30s, and exclusion duration of 90s.

The XCalibur RAW files were collected in profile mode, centroided and converted to MzXML using ReAdW v. 3.5.1. The data was searched using SEQUEST Version 27, rev. 12 (Thermo Fisher Scientific), which was set for two maximum missed cleavages, a precursor mass window of 20ppm, trypsin digestion, variable modification P @ - 27.9949, R @ −43.0534, K @ −1.0316, T @ −2.0157, C @ 57.0293, and M @ 15.9949. Searches were performed with a mouse-specific subset of the UniRefKB database.

#### Peptide Filtering, Grouping, and Quantification

The list of peptide IDs generated based on SEQUEST search results were filtered using Scaffold Version 4.8.9 (Protein Sciences, Portland Oregon). Scaffold filters and groups all peptides to generate and retain only high confidence IDs while also generating normalized spectral counts (N-SC’s) across all samples for the purpose of relative quantification. The filter cut-off values were set with minimum peptide length of >5 AA’s, with no MH+1 charge states, with peptide probabilities of >80% C.I., and with the number of peptides per protein ≥2. The protein probabilities were then set to a >99.0% C.I., and an FDR<1.0. Scaffold incorporates the two most common methods for statistical validation of large proteome datasets, the false discovery rate (FDR) and protein probability (72-74). In addition, for all PTM’s, we further analyzed the exported Scaffold files within Scaffold PTM (Protein Sciences) where the use of assigned A-scores and localization C.I.’s allowed us to filter out potential false positive PTM assignments (75). Those PTM’s that pass these filters were also manually checked for quality of fit.

### Molecular modeling and heme docking

To determine potential mechanisms by which heme exposure impairs ENAC activity within few seconds, we performed computer modeling using YASARA software (35, 36) to simulate heme docking and binding to the known cryo-electron microscopy structure of ENaC (Protein Data Bank: 6BQN) (37). The docking of heme molecule to the ENaC structure was performed using AutoDock program developed at the Scripps Research Institute using the default docking parameters and point charges initially assigned according to the AMBER03 force field (76), and then damped to mimic the less polar Gasteiger charges used to optimize the AutoDock scoring function. The YASARA molecular modeling program was set up to determine the best 25 hits and also determine their free energy of binding in kJ/mol.

## Acknowledgements

The authors would like to thank Dr. Rakesh Patel, Dr. Tamas Jilling for their valuable inputs. In addition, the authors would like to acknowledge technical support of Mark A. Duerr, Jacob D. Franke, and Carolyn J. Albert in the generation of the brominated lipid data.

## Funding

Supported by the CounterACT Program, National Institutes of Health Office of the Director (NIH OD), the National Institute of Neurological Disorders and Stroke (NINDS), and the National Institute of Environmental Health Sciences (NIEHS), Grant Numbers (5UO1 ES026458; 3UO1 ES026458 03S1; 5UO1 ES027697) to **SM** and NIH/NHLBI grant K12 HL143958 to **SA**.

## Notes

**Conflict of interest statement:** The authors have declared that no conflict of interest exists.

## REFERENCES

1. Ryter SW, and Tyrrell RM. The heme synthesis and degradation pathways: role in oxidant sensitivity. Heme oxygenase has both pro- and antioxidant properties. Free radical biology & medicine. 2000;28(2):289–309.

2. Li L, and Frei B. Prolonged exposure to LPS increases iron, heme, and p22phox levels and NADPH oxidase activity in human aortic endothelial cells: inhibition by desferrioxamine. Arterioscler Thromb Vasc Biol. 2009;29(5):732–8.

3. Shannahan JH, Ghio AJ, Schladweiler MC, McGee JK, Richards JH, Gavett SH, and Kodavanti UP. The role of iron in Libby amphibole-induced acute lung injury and inflammation. Inhal Toxicol. 2011;23(6):313–23.

4. Dennery PA, Visner G, Weng Y, Nguyen X, Lu F, Zander D, and Yang G. Resistance to hyperoxia with heme oxygenase-1 disruption: role of iron. Free Radic Biol Med. 2003;34(1):124–33.

5. Kato GJ, Steinberg MH, and Gladwin MT. Intravascular hemolysis and the pathophysiology of sickle cell disease. The Journal of clinical investigation. 2017;127(3):750–60.

6. Stapley R, Rodriguez C, Oh JY, Honavar J, Brandon A, Wagener BM, Marques MB, Weinberg JA, Kerby JD, Pittet JF, et al. Red blood cell washing, nitrite therapy, and antiheme therapies prevent stored red blood cell toxicity after trauma-hemorrhage. Free radical biology & medicine. 2015;85(207-18.

7. Lin T, Maita D, Thundivalappil SR, Riley FE, Hambsch J, Van Marter LJ, Christou HA, Berra L, Fagan S, Christiani DC, et al. Hemopexin in severe inflammation and infection: mouse models and human diseases. Critical care. 2015;19(166.

8. Aggarwal S, Ahmad I, Lam A, Carlisle MA, Li C, Wells JM, Raju SV, Athar M, Rowe SM, Dransfield MT, et al. Heme scavenging reduces pulmonary endoplasmic reticulum stress, fibrosis, and emphysema. JCI insight. 2018;3(21).

9. Aggarwal S, Lam A, Bolisetty S, Carlisle MA, Traylor A, Agarwal A, and Matalon S. Heme Attenuation Ameliorates Irritant Gas Inhalation-Induced Acute Lung Injury. Antioxidants & redox signaling. 2016;24(2):99–112.

10. Aggarwal S, Jilling T, Doran S, Ahmad I, Eagen JE, Gu S, Gillespie M, Albert CJ, Ford D, Oh JY, et al. Phosgene inhalation causes hemolysis and acute lung injury. Toxicology letters. 2019;312(204-13.

11. Higdon AN, Benavides GA, Chacko BK, Ouyang X, Johnson MS, Landar A, Zhang J, and Darley-Usmar VM. Hemin causes mitochondrial dysfunction in endothelial cells through promoting lipid peroxidation: the protective role of autophagy. Am J Physiol Heart Circ Physiol. 2012;302(7):H1394–409.

12. Gaggar A, and Patel RP. There is blood in the water: Hemolysis, hemoglobin, and heme in acute lung injury. Am J Physiol Lung Cell Mol Physiol. 2016:ajplung 00312 2016.

13. Suliman HB, Carraway MS, Velsor LW, Day BJ, Ghio AJ, and Piantadosi CA. Rapid mtDNA deletion by oxidants in rat liver mitochondria after hemin exposure. Free radical biology & medicine. 2002;32(3):246–56.

14. Fortes GB, Alves LS, de Oliveira R, Dutra FF, Rodrigues D, Fernandez PL, Souto-Padron T, De Rosa MJ, Kelliher M, Golenbock D, et al. Heme induces programmed necrosis on macrophages through autocrine TNF and ROS production. Blood. 2012;119(10):2368–75.

15. Balla J, Vercellotti GM, Jeney V, Yachie A, Varga Z, Jacob HS, Eaton JW, and Balla G. Heme, heme oxygenase, and ferritin: how the vascular endothelium survives (and dies) in an iron-rich environment. Antioxid Redox Signal. 2007;9(12):2119–37.

16. Heide K, Haupt H, Stoeriko K, and Schultze HE. On the Heme-Binding Capacity of Hemopexin. Clin Chim Acta. 1964;10(460-9.

17. Killander J. Separation of Human Heme- and Hemoglobin-Binding Plasma Proteins, Ceruloplasmin and Albumin by Gel Filtration. Biochim Biophys Acta. 1964;93(1-14.

18. Miller YI, Smith A, Morgan WT, and Shaklai N. Role of hemopexin in protection of low-density lipoprotein against hemoglobin-induced oxidation. Biochemistry. 1996;35(40):13112–7.

19. Nagel RL, and Gibson QH. The binding of hemoglobin to haptoglobin and its relation to subunit dissociation of hemoglobin. J Biol Chem. 1971;246(1):69–73.

20. Shaver CM, Upchurch CP, Janz DR, Grove BS, Putz ND, Wickersham NE, Dikalov SI, Ware LB, and Bastarache JA. Cell-free hemoglobin: a novel mediator of acute lung injury. American journal of physiology Lung cellular and molecular physiology. 2016;310(6):L532–41.

21. Matalon S. Mechanisms and regulation of ion transport in adult mammalian alveolar type II pneumocytes. The American journal of physiology. 1991;261(5 Pt 1):C727–38.

22. Ware LB, and Matthay MA. Alveolar fluid clearance is impaired in the majority of patients with acute lung injury and the acute respiratory distress syndrome. American journal of respiratory and critical care medicine. 2001;163(6):1376–83.

23. Vivona ML, Matthay M, Chabaud MB, Friedlander G, and Clerici C. Hypoxia reduces alveolar epithelial sodium and fluid transport in rats: reversal by beta-adrenergic agonist treatment. American journal of respiratory cell and molecular biology. 2001;25(5):554–61.

24. Gwozdzinska P, Buchbinder BA, Mayer K, Herold S, Morty RE, Seeger W, and Vadasz I. Hypercapnia Impairs ENaC Cell Surface Stability by Promoting Phosphorylation, Polyubiquitination and Endocytosis of beta-ENaC in a Human Alveolar Epithelial Cell Line. Frontiers in immunology. 2017;8(591.

25. Lam A, Vetal N, Matalon S, and Aggarwal S. Role of heme in bromine-induced lung injury. Annals of the New York Academy of Sciences. 2016;1374(1):105–10.

26. Ford DA, Honavar J, Albert CJ, Duerr MA, Oh JY, Doran S, Matalon S, and Patel RP. Formation of chlorinated lipids post-chlorine gas exposure. Journal of lipid research. 2016;57(8):1529–40.

27. Lazrak A, Creighton J, Yu Z, Komarova S, Doran SF, Aggarwal S, Emala CW, Sr., Stober VP, Trempus CS, Garantziotis S, et al. Hyaluronan mediates airway hyperresponsiveness in oxidative lung injury. American journal of physiology Lung cellular and molecular physiology. 2015;308(9):L891–903.

28. Meyer NJ, Reilly JP, Feng R, Christie JD, Hazen SL, Albert CJ, Franke JD, Hartman CL, McHowat J, and Ford DA. Myeloperoxidase-derived 2-chlorofatty acids contribute to human sepsis mortality via acute respiratory distress syndrome. JCI insight. 2017;2(23).

29. Duerr MA, Aurora R, and Ford DA. Identification of glutathione adducts of alpha-chlorofatty aldehydes produced in activated neutrophils. Journal of lipid research. 2015;56(5):1014–24.

30. Duerr MA, Palladino END, Hartman CL, Lambert JA, Franke JD, Albert CJ, Matalon S, Patel RP, Slungaard A, and Ford DA. Bromofatty aldehyde derived from bromine exposure and myeloperoxidase and eosinophil peroxidase modify GSH and protein. Journal of lipid research. 2018;59(4):696–705.

31. Chen L, Patel RP, Teng X, Bosworth CA, Lancaster JR, Jr., and Matalon S. Mechanisms of cystic fibrosis transmembrane conductance regulator activation by S-nitrosoglutathione. The Journal of biological chemistry. 2006;281(14):9190–9.

32. Guo Y, DuVall MD, Crow JP, and Matalon S. Nitric oxide inhibits Na+ absorption across cultured alveolar type II monolayers. The American journal of physiology. 1998;274(3):L369–77.

33. Lazrak A, Iles KE, Liu G, Noah DL, Noah JW, and Matalon S. Influenza virus M2 protein inhibits epithelial sodium channels by increasing reactive oxygen species. FASEB journal : official publication of the Federation of American Societies for Experimental Biology. 2009;23(11):3829–42.

34. Lazrak A, Chen L, Jurkuvenaite A, Doran SF, Liu G, Li Q, Lancaster JR, Jr., and Matalon S. Regulation of alveolar epithelial Na+ channels by ERK1/2 in chlorine-breathing mice. American journal of respiratory cell and molecular biology. 2012;46(3):342–54.

35. Krieger E, Koraimann G, and Vriend G. Increasing the precision of comparative models with YASARA NOVA--a self-parameterizing force field. Proteins. 2002;47(3):393–402.

36. Aggarwal S, Gross CM, Rafikov R, Kumar S, Fineman JR, Ludewig B, Jonigk D, and Black SM. Nitration of tyrosine 247 inhibits protein kinase G-1alpha activity by attenuating cyclic guanosine monophosphate binding. The Journal of biological chemistry. 2014;289(11):7948–61.

37. Noreng S, Bharadwaj A, Posert R, Yoshioka C, and Baconguis I. Structure of the human epithelial sodium channel by cryo-electron microscopy. eLife. 2018;7(

38. Gessner MA, Doran SF, Yu Z, Dunaway CW, Matalon S, and Steele C. Chlorine gas exposure increases susceptibility to invasive lung fungal infection. American journal of physiology Lung cellular and molecular physiology. 2013;304(11):L765–73.

39. Honavar J, Doran S, Ricart K, Matalon S, and Patel RP. Nitrite therapy prevents chlorine gas toxicity in rabbits. Toxicology letters. 2017;271(20-5.

40. Wagener FA, Eggert A, Boerman OC, Oyen WJ, Verhofstad A, Abraham NG, Adema G, van Kooyk Y, de Witte T, and Figdor CG. Heme is a potent inducer of inflammation in mice and is counteracted by heme oxygenase. Blood. 2001;98(6):1802–11.

41. Muller-Eberhard U, and Fraig M. Bioactivity of heme and its containment. American journal of hematology. 1993;42(1):59–62.

42. Reiter CD, Wang X, Tanus-Santos JE, Hogg N, Cannon RO, 3rd, Schechter AN, and Gladwin MT. Cell-free hemoglobin limits nitric oxide bioavailability in sickle-cell disease. Nature medicine. 2002;8(12):1383–9.

43. Squadrito GL, Postlethwait EM, and Matalon S. Elucidating mechanisms of chlorine toxicity: reaction kinetics, thermodynamics, and physiological implications. American journal of physiology Lung cellular and molecular physiology. 2010;299(3):L289–300.

44. Chou AC, and Fitch CD. Mechanism of hemolysis induced by ferriprotoporphyrin IX. The Journal of clinical investigation. 1981;68(3):672–7.

45. Kirschner-Zilber I, Rabizadeh E, and Shaklai N. The interaction of hemin and bilirubin with the human red cell membrane. Biochimica et biophysica acta. 1982;690(1):20–30.

46. Liu SC, Zhai S, Lawler J, and Palek J. Hemin-mediated dissociation of erythrocyte membrane skeletal proteins. The Journal of biological chemistry. 1985;260(22):12234–9.

47. Solar I, Dulitzky J, and Shaklai N. Hemin-promoted peroxidation of red cell cytoskeletal proteins. Archives of biochemistry and biophysics. 1990;283(1):81–9.

48. Szweda-Lewandowska Z, Krokosz A, Gonciarz M, Zajeczkowska W, and Puchala M. Damage to human erythrocytes by radiation-generated HO* radicals: molecular changes in erythrocyte membranes. Free radical research. 2003;37(10):1137–43.

49. Matthay MA, Folkesson HG, and Clerici C. Lung epithelial fluid transport and the resolution of pulmonary edema. Physiological reviews. 2002;82(3):569–600.

50. Mutlu GM, and Sznajder JI. Mechanisms of pulmonary edema clearance. American journal of physiology Lung cellular and molecular physiology. 2005;289(5):L685–95.

51. Matalon S, and O’Brodovich H. Sodium channels in alveolar epithelial cells: molecular characterization, biophysical properties, and physiological significance. Annual review of physiology. 1999;61(627-61.

52. Hummler E, Barker P, Gatzy J, Beermann F, Verdumo C, Schmidt A, Boucher R, and Rossier BC. Early death due to defective neonatal lung liquid clearance in alpha-ENaC-deficient mice. Nature genetics. 1996;12(3):325–8.

53. Thome U, Chen L, Factor P, Dumasius V, Freeman B, Sznajder JI, and Matalon S. Na,K-ATPase gene transfer mitigates an oxidant-induced decrease of active sodium transport in rat fetal ATII cells. American journal of respiratory cell and molecular biology. 2001;24(3):245–52.

54. Brand JD, Lazrak A, Trombley JE, Shei RJ, Adewale AT, Tipper JL, Yu Z, Ashtekar AR, Rowe SM, Matalon S, et al. Influenza-mediated reduction of lung epithelial ion channel activity leads to dysregulated pulmonary fluid homeostasis. JCI insight. 2018;3(20).

55. Brazee PL, Morales-Nebreda L, Magnani ND, Garcia JG, Misharin AV, Ridge KM, Budinger GRS, Iwai K, Dada LA, and Sznajder JI. Linear ubiquitin assembly complex regulates lung epithelial driven responses during influenza infection. The Journal of clinical investigation. 2019.

56. Factor P, Saldias F, Ridge K, Dumasius V, Zabner J, Jaffe HA, Blanco G, Barnard M, Mercer R, Perrin R, et al. Augmentation of lung liquid clearance via adenovirus-mediated transfer of a Na,K-ATPase beta1 subunit gene. The Journal of clinical investigation. 1998;102(7):1421–30.

57. Peteranderl C, Morales-Nebreda L, Selvakumar B, Lecuona E, Vadasz I, Morty RE, Schmoldt C, Bespalowa J, Wolff T, Pleschka S, et al. Macrophage-epithelial paracrine crosstalk inhibits lung edema clearance during influenza infection. The Journal of clinical investigation. 2016;126(4):1566–80.

58. Ji HL, Su XF, Kedar S, Li J, Barbry P, Smith PR, Matalon S, and Benos DJ. Delta-subunit confers novel biophysical features to alpha beta gamma-human epithelial sodium channel (ENaC) via a physical interaction. The Journal of biological chemistry. 2006;281(12):8233–41.

59. Otulakowski G, Rafii B, and O’Brodovich H. Differential translational efficiency of ENaC subunits during lung development. American journal of respiratory cell and molecular biology. 2004;30(6):862–70.

60. Frindt G, Ergonul Z, and Palmer LG. Na channel expression and activity in the medullary collecting duct of rat kidney. American journal of physiology Renal physiology. 2007;292(4):F1190–6.

61. Ismailov, II, Berdiev BK, and Benos DJ. Biochemical status of renal epithelial Na+ channels determines apparent channel conductance, ion selectivity, and amiloride sensitivity. Biophysical journal. 1995;69(5):1789–800.

62. Butterworth MB, Edinger RS, Frizzell RA, and Johnson JP. Regulation of the epithelial sodium channel by membrane trafficking. American journal of physiology Renal physiology. 2009;296(1):F10–24.

63. Butterworth MB, Edinger RS, Johnson JP, and Frizzell RA. Acute ENaC stimulation by cAMP in a kidney cell line is mediated by exocytic insertion from a recycling channel pool. The Journal of general physiology. 2005;125(1):81–101.

64. Butterworth MB, Helman SI, and Els WJ. cAMP-sensitive endocytic trafficking in A6 epithelia. American journal of physiology Cell physiology. 2001;280(4):C752–62.

65. Wang S, Publicover S, and Gu Y. An oxygen-sensitive mechanism in regulation of epithelial sodium channel. Proceedings of the National Academy of Sciences of the United States of America. 2009;106(8):2957–62.

66. Tang XD, Xu R, Reynolds MF, Garcia ML, Heinemann SH, and Hoshi T. Haem can bind to and inhibit mammalian calcium-dependent Slo1 BK channels. Nature. 2003;425(6957):531–5.

67. Albert CJ, Thukkani AK, Heuertz RM, Slungaard A, Hazen SL, and Ford DA. Eosinophil peroxidase-derived reactive brominating species target the vinyl ether bond of plasmalogens generating a novel chemoattractant, alpha-bromo fatty aldehyde. The Journal of biological chemistry. 2003;278(11):8942–50.

68. Wacker BK, Albert CJ, Ford BA, and Ford DA. Strategies for the analysis of chlorinated lipids in biological systems. Free Radic Biol Med. 2013;59(92-9.

69. Pan D, Vargas-Morales O, Zern B, Anselmo AC, Gupta V, Zakrewsky M, Mitragotri S, and Muzykantov V. The Effect of Polymeric Nanoparticles on Biocompatibility of Carrier Red Blood Cells. PLoS One. 2016;11(3):e0152074.

70. Lazrak A, Jurkuvenaite A, Chen L, Keeling KM, Collawn JF, Bedwell DM, and Matalon S. Enhancement of alveolar epithelial sodium channel activity with decreased cystic fibrosis transmembrane conductance regulator expression in mouse lung. American journal of physiology Lung cellular and molecular physiology. 2011;301(4):L557–67.

71. Lazrak A, Nita I, Subramaniyam D, Wei S, Song W, Ji HL, Janciauskiene S, and Matalon S. Alpha(1)-antitrypsin inhibits epithelial Na+ transport in vitro and in vivo. American journal of respiratory cell and molecular biology. 2009;41(3):261–70.

72. Keller A, Nesvizhskii AI, Kolker E, and Aebersold R. Empirical statistical model to estimate the accuracy of peptide identifications made by MS/MS and database search. Analytical chemistry. 2002;74(20):5383–92.

73. Nesvizhskii AI, Keller A, Kolker E, and Aebersold R. A statistical model for identifying proteins by tandem mass spectrometry. Analytical chemistry. 2003;75(17):4646–58.

74. Weatherly DB, Atwood JA, 3rd, Minning TA, Cavola C, Tarleton RL, and Orlando R. A Heuristic method for assigning a false-discovery rate for protein identifications from Mascot database search results. Molecular & cellular proteomics : MCP. 2005;4(6):762–72.

75. Beausoleil SA, Villen J, Gerber SA, Rush J, and Gygi SP. A probability-based approach for high-throughput protein phosphorylation analysis and site localization. Nature biotechnology. 2006;24(10):1285–92.

76. Duan Y, Wu C, Chowdhury S, Lee MC, Xiong G, Zhang W, Yang R, Cieplak P, Luo R, Lee T, et al. A point-charge force field for molecular mechanics simulations of proteins based on condensed-phase quantum mechanical calculations. Journal of computational chemistry. 2003;24(16):1999–2012.

