## Supplemental Figures 1,2,3 for "Heme Impairs Alveolar Epithelial Sodium Channels Post Toxic Gas Inhalation"

**Supplemental Figures and Legends**

**Supplement Figure 1. Brominated fatty aldehyde (BrFALD) and fatty acids (BrFA) levels in broncholaveolar lavage fluid (BALF) of Br_2_ exposed mice.** Adult male C57BL/6 mice were exposed to Br_2_ gas (600ppm, 30min) and returned them to room air. Using ESI-LC/MS/MS quantitation, we measured 16- and 18- carbon BrFALD and BrFA in the BALF of mice at different intervals post exposure. The levels of 16BrFALD (n=6-13) (A), total 16BrFA (n=6-13) (B), free 16BrFA (n=6-13) (C), and esterified 16BrFA (n=6-13) (D) increased in BALF of Br_2_ exposed mice. Similarly, the levels of 18BrFALD (n=6-13) (E), total 18BrFA (n=6-13) (F), free 18BrFA (n=6-13) (G), and esterified 18BrFA (n=6-13) (H) increased in the BALF of Br_2_ exposed mice. Values and means ± SEM. **P* < 0.05 vs. air exposed mice, by one-way ANOVA followed by Tukey post hoc testing.

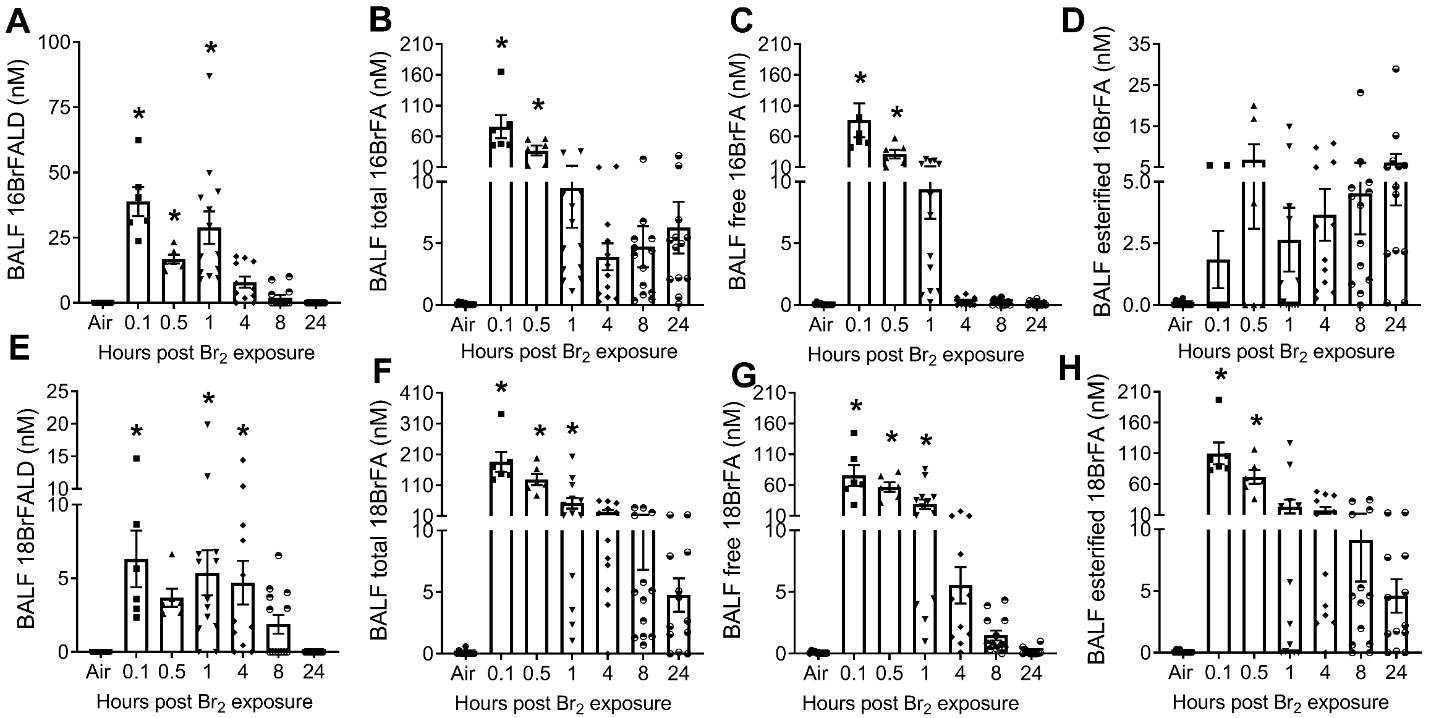

**Supplement Figure 2. Glutathionylated fatty aldehyde (FALD-GSH) levels in broncholaveolar lavage fluid (BALF) and peripheral lung tissue of Br_2_ exposed mice.** Adult male C57BL/6 mice were exposed to Br_2_ gas (600ppm, 30min) and returned them to room air. Using ESI-LC/MS/MS quantitation, we measured 16- and 18- carbon BrFALD-GSH in the BALF and the peripheral lung tissue of mice at different intervals post exposure. The levels of 16FALD-GSH (n=6-13) (A) and 18FALD-GSH (n=6-13) (B) increased in BALF of Br_2_ exposed mice. Similarly, the levels of 16FALD-GSH (n=6-13) (C) and 18FALD-GSH (n=6-13) (D) increased in the peripheral lung tissue of Br_2_ exposed mice. Values are means ± SEM. **P* < 0.05 vs. air exposed mice, by one-way ANOVA followed by Tukey post hoc testing.

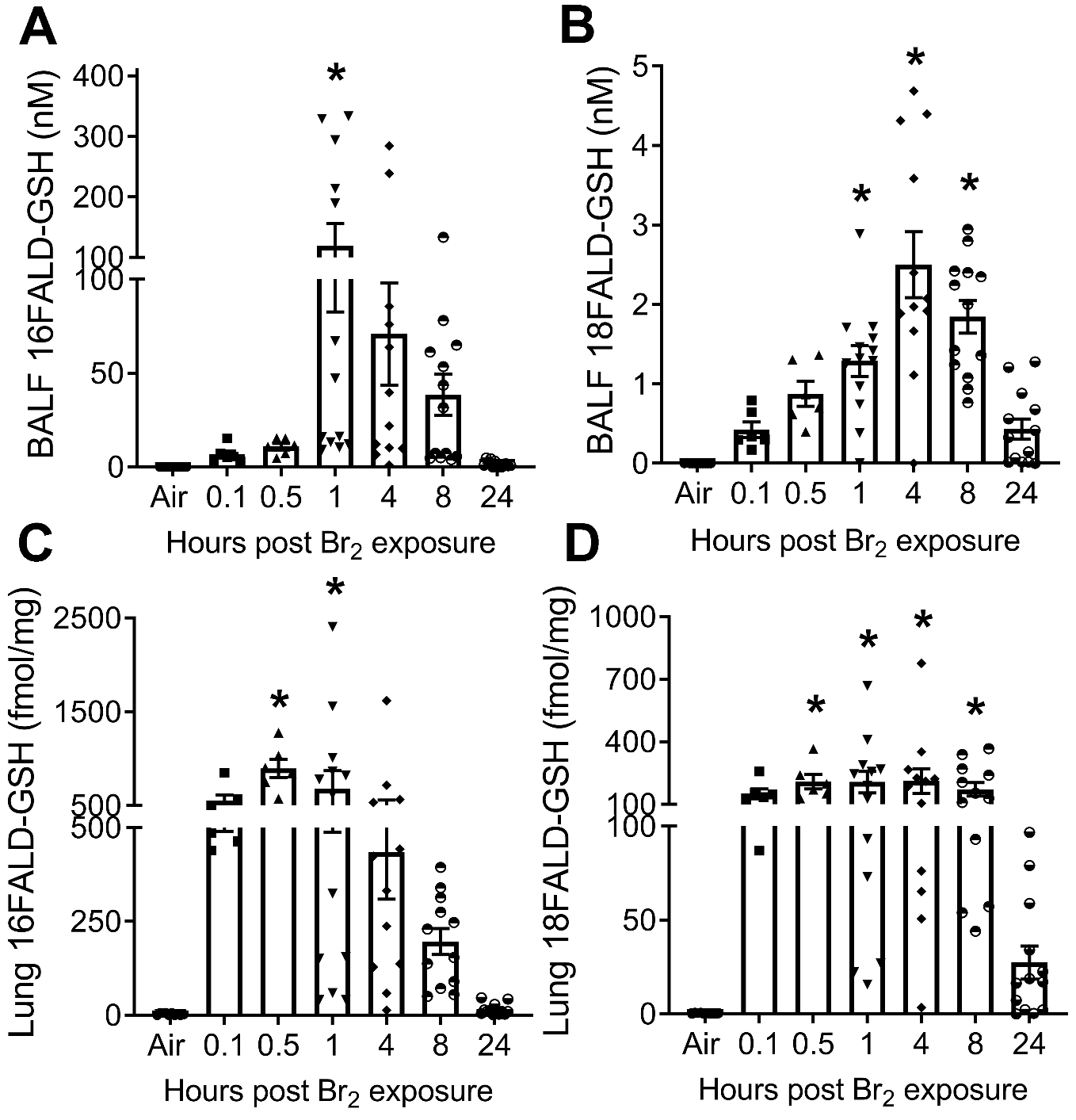

**Supplement Figure 3. Glutathionylated fatty aldehyde (FALD-GSH) levels in plasma, urine, and RBCs of Br_2_ exposed mice.** Adult male C57BL/6 mice were exposed to Br_2_ gas (600ppm, 30min) and returned them to room air. Using ESI-LC/MS/MS quantitation, we measured 16- and 18- carbon FALD-GSH in plasma, urine, and RBCs of mice at different intervals post exposure. The levels of 16FALD-GSH (n=6-13) (A) and 18FALD-GSH (n=6-13) (B) increased in plasma of Br_2_ exposed mice. The levels of 16FALD-GSH (n=4-5) (C) and 18FALD-GSH (n=4-5) (D) also increased in the urine of Br_2_ exposed mice. Similarly, the levels of 16FALD-GSH (n=4-5) (C) and 18FALD-GSH (n=4-5) (D) were elevated in the RBCs of Br_2_ exposed mice. Values are means ± SEM. **P* < 0.05 vs. air exposed mice, by one-way ANOVA followed by Tukey post hoc testing.

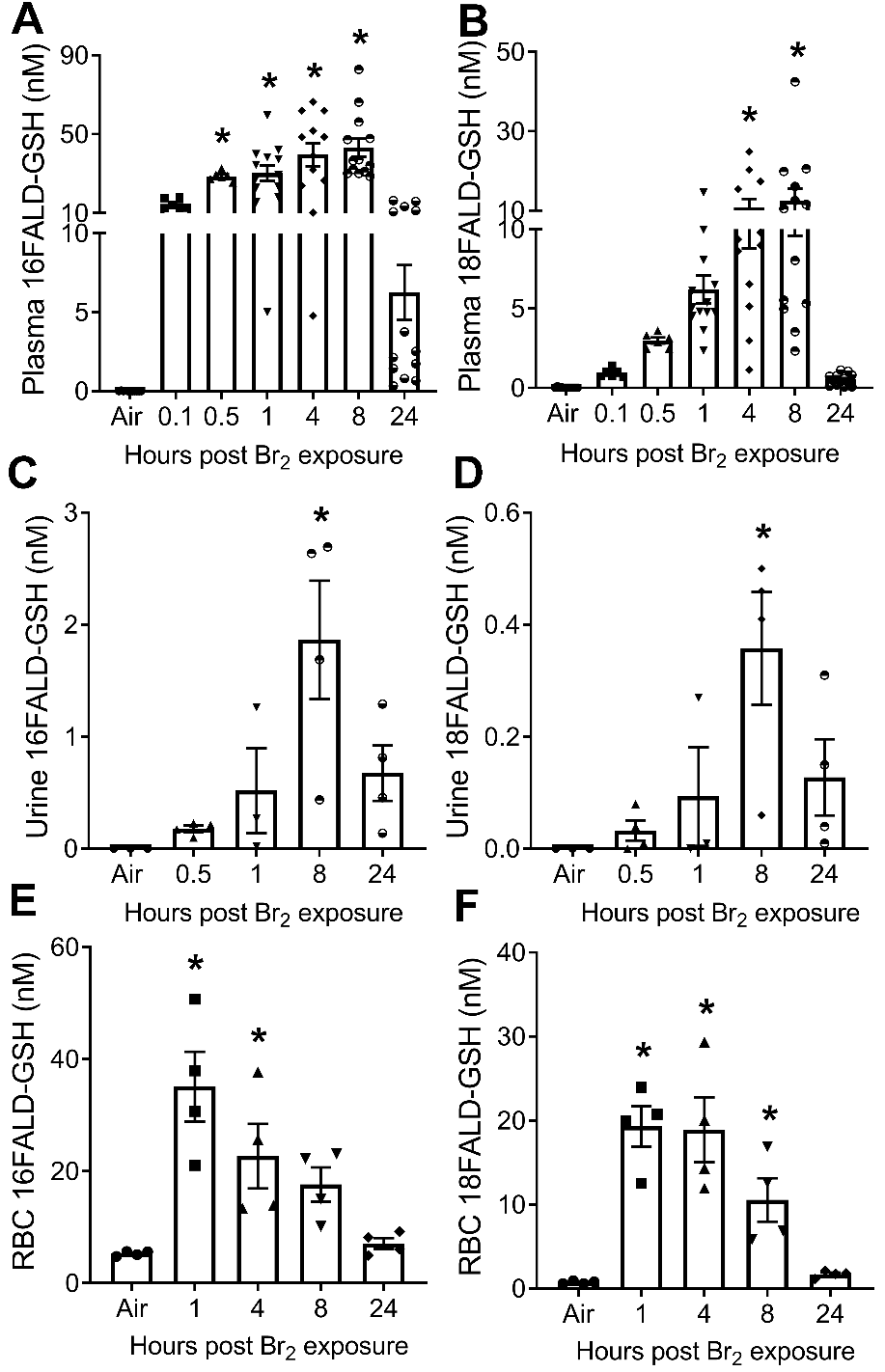
